# Apoplast-localized β-Glucosidase Elevates Isoflavone Accumulation in the Soybean Rhizosphere

**DOI:** 10.1101/2022.12.13.520340

**Authors:** Hinako Matsuda, Yumi Yamazaki, Eiko Moriyoshi, Masaru Nakayasu, Shinichi Yamazaki, Yuichi Aoki, Hisabumi Takase, Shin Okazaki, Atsushi J. Nagano, Akito Kaga, Kazufumi Yazaki, Akifumi Sugiyama

## Abstract

Plant specialized metabolites (PSMs) are often stored as glycosides within cells and released from the roots with some chemical modifications. While isoflavones are known to function as symbiotic signals with rhizobia and to modulate the soybean rhizosphere microbiome, the underlying mechanisms of root-to-soil delivery are poorly understood. In addition to transporter-mediated secretion, the hydrolysis of isoflavone glycosides in the apoplast by an isoflavone conjugate-hydrolyzing β-glucosidase (ICHG) has been proposed but not yet verified. To clarify the role of ICHG in isoflavone supply to the rhizosphere, we have isolated two independent mutants defective in ICHG activity from a soybean high-density mutant library. In the *ichg* mutants, the isoflavone contents and composition in the root apoplast and root exudate significantly changed. When grown in a field, the lack of ICHG activity considerably reduced isoflavone aglycone contents in roots and the rhizosphere soil, although the transcriptomes showed no distinct differences between the *ichg* mutants and WTs. Despite the change in isoflavone contents and composition of the root and rhizosphere of the mutants, root and rhizosphere bacterial communities were not distinctive from those of the WTs. Root bacterial communities and nodulation capacities of the *ichg* mutants did not differ from the WTs under nitrogen-deficient conditions, either. Taken together, these results indicate that ICHG elevates the accumulation of isoflavones in the soybean rhizosphere but is not essential in isoflavone-mediated plant-microbe interactions.

## Introduction

It is predicted that plant species produce over one million different organic compounds (Afendi et al., 2012). It has also been found that approximately 30–60 % of net fixed carbon is transferred to plant roots and approximately 40–90 % of this transferred carbon is released into the rhizosphere, depending on the plant species, age, and environmental conditions (Lynch and Whipps, 1990). Root exudates contain amino acids, sugars, organic acids, fatty acids, vitamins, nucleotides, and specialized (secondary) metabolites (Vives-Peris et al., 2020), and these plant-borne metabolites affect rhizosphere soil properties, such as aggregation, pH, and the microbiome (Ehrenfeld et al., 2005). Plant specialized metabolites (PSMs) have gained attention as they are critical for improving plant nutrition, repelling pathogens and pests, and modulating abiotic stress tolerance (Jacoby et al., 2021; Massalha et al., 2017; Pang et al., 2021; Pascale et al., 2019). Recent studies have demonstrated that PSMs modulate the composition and function of the root and rhizosphere microbiome, influencing both plant growth and fitness. Flavonoids, one of the common PSMs in the rhizosphere, function as signaling molecules to induce rhizobial Nod factors and initiate legume-rhizobia symbiosis (Subramanian et al., 2007), and as iron chelators to promote iron availability (Cesco et al., 2010). Flavonoids can also enrich specific bacterial families in the rhizosphere microbiota, such as *Comamonadaceae* in soybean and *Oxalobacteraceae* in maize, improving maize performance in nitrogen-deficient conditions (Okutani et al., 2020; Yu et al., 2021).

Once flavonoids are synthesized at the surface of the endoplasmic reticulum in the cytosol, they are transported into the vacuole for storage (Marrs et al., 1995; Zhao and Dixon, 2010). Due to the toxicity, instability, and/or insolubility of aglycones in the cells, flavonoids exist as glycosides in the vacuole (Le Roy et al., 2016; Matern et al., 1983). β-Glucosidases, often found in the plastid and apoplast, hydrolyze a variety of glycosides into aglycones when glycosides are delivered from the vacuole for defense or the development of symbiotic relationships (Ketudat Cairns and Esen, 2010; Morant et al., 2008). In soybeans, isoflavone aglycones such as daidzein and genistein serve as initiation signals for symbiosis with *Bradyrhizobium diazoefficiens* and *Ensifer fredii* (Kosslak et al., 1987; Pueppke et al., 1998), and also affect the assembly of the rhizosphere microbiome (Okutani et al., 2020).

Both plasma membrane-localized ATP-binding cassette (ABC) transporters and multidrug and toxic compound extrusion (MATE) transporters mediate the secretion of (iso)flavonoid aglycones to the rhizosphere (Biala et al., 2017; Biała-Leonhard et al., 2021; Sugiyama et al., 2007). In addition, isoflavone glycosides stored in the vacuole are proposed to be secreted to the apoplast, since isoflavone conjugate-hydrolyzing β-glucosidase (ICHG) hydrolyzes these glycosides to aglycones in the apoplast (Fig. 1) (Suzuki et al., 2006).

**Figure 1.**
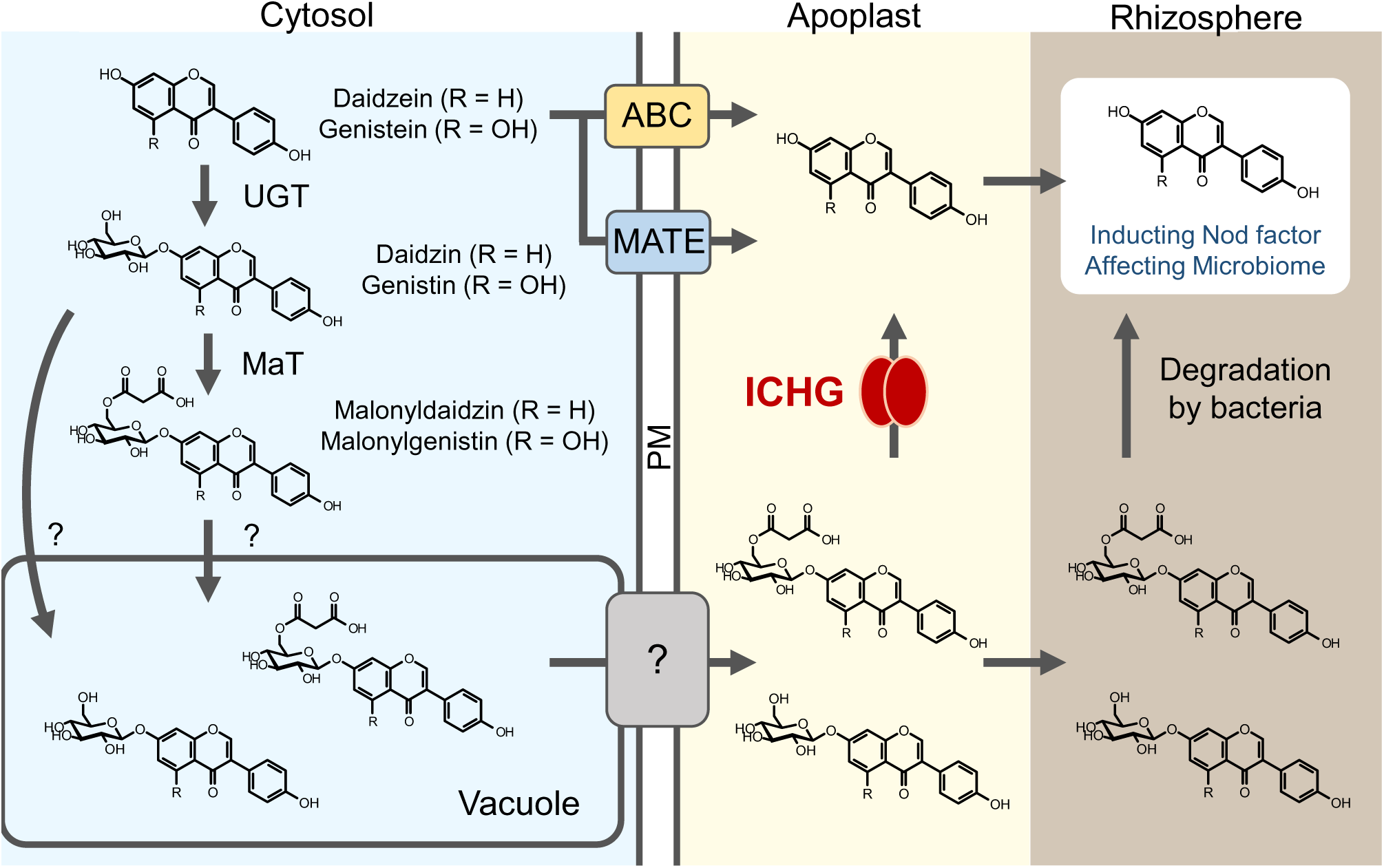
Proposed isoflavone supply routes from the cytosol to the rhizosphere. The apoplast includes the cell walls and intercellular space. ABC, ATP-binding cassette transporter; ICHG, isoflavone conjugate-hydrolyzing β-glucosidase; MaT, malonyl transferase; MATE, multidrug and toxic compound extrusion transporter; PM, plasma membrane; UGT, UDP-glucuronosyltransferase.

ICHG is expressed in the roots, especially from the stele to lateral root hairs, and localized in the cell walls and intercellular space (Suzuki et al., 2006; Yoo et al., 2013). ICHG efficiently hydrolyzes isoflavone malonylglucosides and glucosides (Suzuki et al., 2006). Previous studies have suggested that there is a relationship between *ICHG* gene expression and isoflavone secretion during the different developmental stages of soybean and its diurnal regulation when hydroponically grown. From the vegetative to reproductive stages, *ICHG* gene expression levels were found to decrease while the isoflavone glycoside contents in the hydroponic culture increased (Sugiyama et al., 2016). The expression levels of the isoflavone biosynthetic genes and candidate isoflavone transporter genes increased around noon each day and then decreased around midnight, but the isoflavone aglycone content in the hydroponic medium remained constant (Matsuda et al., 2020). The gene expression of *ICHG* exhibited the opposite diurnal pattern against the isoflavone biosynthetic genes and candidate transporters, suggesting that ICHG degrades the glycosides secreted from the vacuole to complement the decreased transporter-mediated secretion of isoflavone aglycones during the night (Matsuda et al., 2020). Despite these gene expression analyses and enzymatic characterizations, the physiological function of ICHG remains unknown. In this study, we employed soybean mutants defective in ICHG to clarify how ICHG contributes to the delivery of isoflavones from soybean roots to the rhizosphere and their interactions with soil bacteria.

## Results

### Development of ichg mutants

We screened *ichg* mutants from a high-density mutant library, generated using an ethyl methanesulfonate (EMS) treatment twice, via an amplicon sequencing method (Tsuda et al., 2015). Two *ichg* mutants, with missense (Glu420Lys hereafter *ichg-1*) and nonsense (Gln232stop hereafter *ichg-2*) mutations, respectively, in the *ICHG* gene (*Glyma.12G053800*), were identified from the 1,536 mutants in the library (Fig. 2A). These mutants were backcrossed with the wild-type cultivar Enrei, F_1_ plants were subjected to self-pollination, and seeds obtained from the F_2_ plants (the missense mutant-derived homozygous *ichg-1* and homozygous wild-type (WT)-1 plants; the nonsense mutant-derived homozygous *ichg-2* and homozygous WT-2 plants) were used in the following experiments (Fig. 2B).

**Figure 2.**
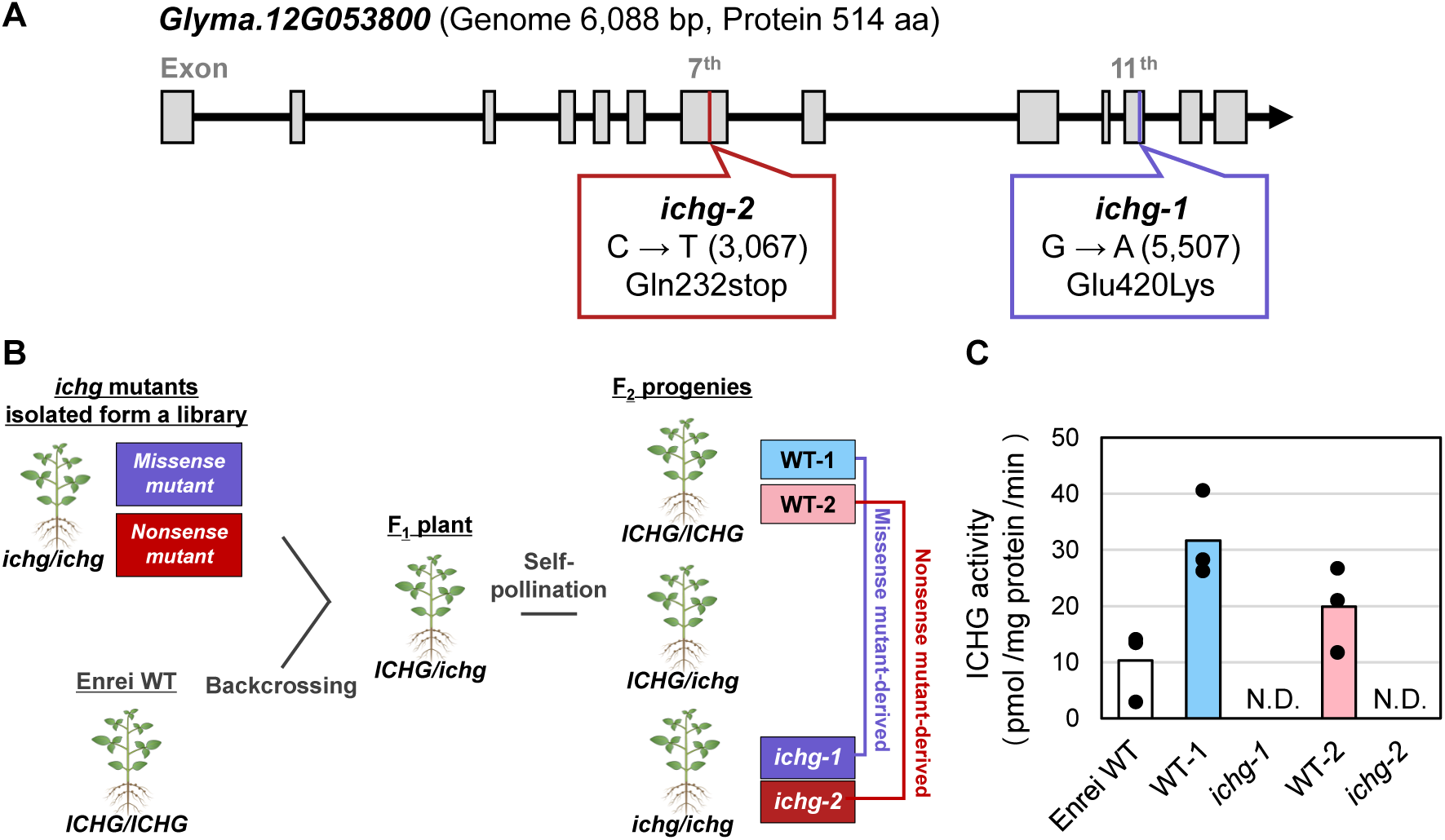
Development of *ichg* mutants from a soybean high-density mutant library. (A) Gene mutations and amino acid replacements in the *ICHG* gene of *ichg-1* and *ichg-2* (*Glyma.12G053800*, genome sequence 6,088 bp from start to stop codon, peptide sequence 514 aa). Gray squares show exons of *ICHG*. In *ichg-1*, the 420^th^ glutamic acid is replaced by lysine (peptide sequence 514 aa). In *ichg-2*, the 232^nd^ glutamine is replaced by a stop codon (peptide sequence 231 aa). (B) Scheme for acquiring homozygous *ichg* mutants and WT alleles. Missense and nonsense *ichg* mutants from the soybean (cultivar Enrei) mutant library were backcrossed with Enrei WT, and F_2_ progenies were obtained through the self-pollination of the F_1_ plants. (C) ICHG activity of the apoplastic crude proteins extracted from the roots of WT soybean and two homozygous *ichg* mutants and their WT alleles. The individual black dots indicate raw data points. The bar graphs show the mean of three technical replicates. ICHG activity of the apoplastic crude proteins extracted from *ichg-1* and *ichg-2* was below the detection limit. ICHG, isoflavone conjugate-hydrolyzing β-glucosidase; N.D., not detected; WT, wild-type.

Heterologously expressed ICHG in *Escherichia coli* specifically degrades isoflavone glycosides, and malonylglucoside has been identified as the best substrate (Suzuki et al., 2006). We evaluated malonyldaidzin degradation activity using apoplastic crude proteins extracted from the roots of the F_2_ progenies. The ICHG activity in the crude protein from the soybeans with either the *ichg-1* or *ichg-2* alleles was below the detection limit (Fig. 2C), while that of the WTs was apparent. These results confirmed that the activity of ICHG in the apoplastic crude proteins extracted from both *ichg* mutants was completely lost, and thus, these mutants could be utilized in further analyses.

### Isoflavone contents in the root apoplastic fraction and hydroponic medium of ichg mutants

To verify the effects of the ICHG defects in the root apoplast where ICHG is localized, we analyzed isoflavone aglycones and glycosides in the root apoplastic fraction collected from 1-week-old soybean seedlings. The amount of daidzein, the most abundant isoflavone aglycone in soybean roots, was much lower in the apoplastic fraction of *ichg-1* and *ichg-2* than the WTs (*p* = 0.008) (Fig. 3A). Daidzein levels in the whole roots of *ichg-1* and *ichg-2* were also lower than those of the WTs (*ichg-1*, *p* = 0.009; *ichg-2*, *p* = 0.035) (Fig. 3C). The amount of genistein, the next abundant aglycone, in the apoplastic fraction and roots of the *ichg* mutants showed not a significant but declining trend when compared with the WTs (apoplastic fluids: *ichg-1*, *p* = 0.178; *ichg-2*, *p* = 0.103; whole roots: *ichg-1*, *p* = 0.006; *ichg-2*, *p* = 0.020) (Fig. 3A, C). The contents of isoflavone glycosides including daidzin, genistin, malonyldaidzin, and malonylgenistin were significantly higher in the root apoplastic fractions of the *ichg* mutants than those of the WTs, while their contents in the roots were comparable between the *ichg* mutants and WTs (Fig. 3A, C). In addition, the total isoflavone contents in the apoplastic fractions were also considerably higher in the *ichg* mutants than the WTs (*ichg-1*, *p* = 0.062; *ichg-2*, *p* = 0.049) (Fig. 3B); while no clear difference was observed in the total isoflavone content of the whole roots (Fig. 3D).

**Figure 3.**
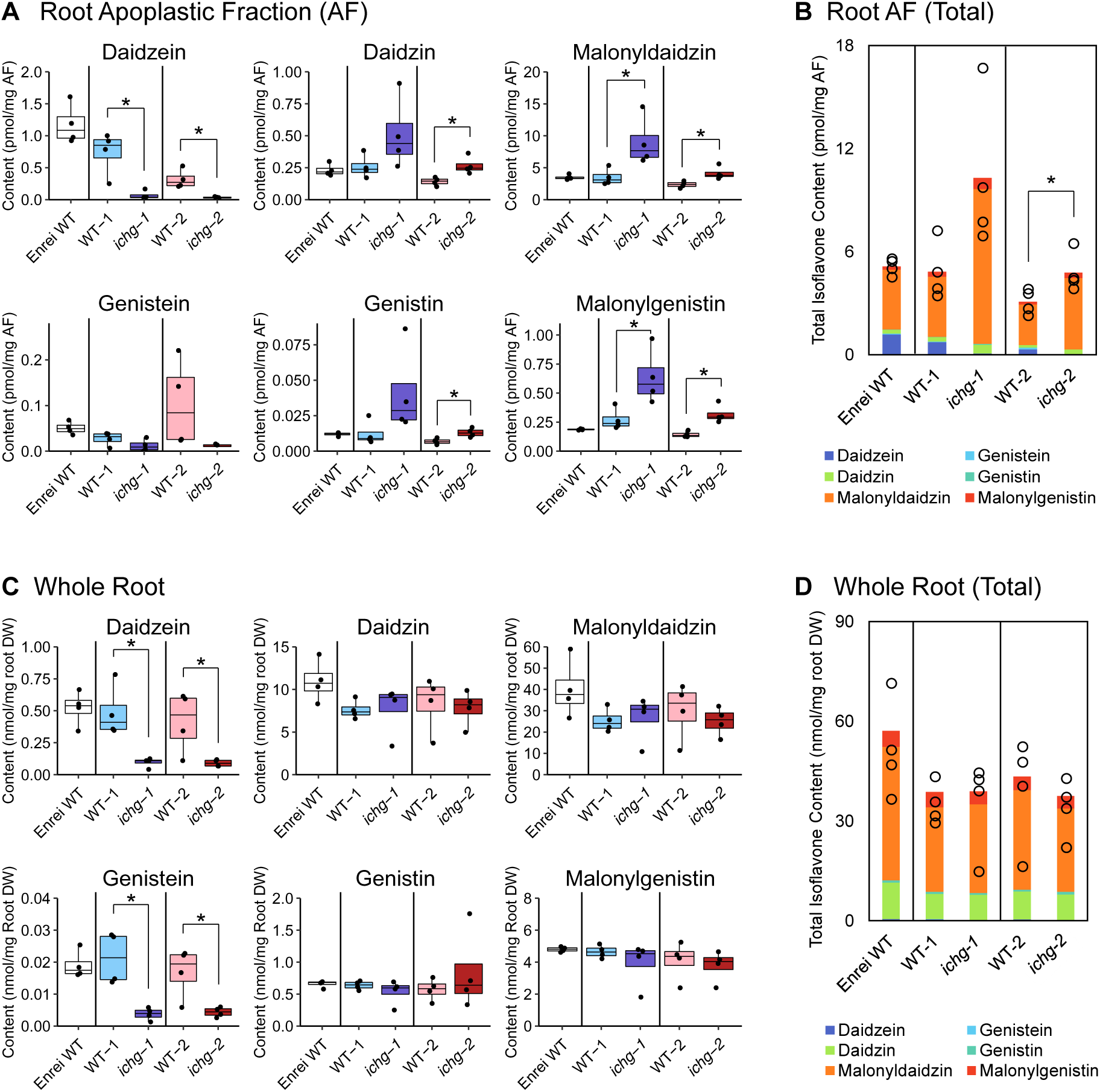
Individual and total contents of the major isoflavones in the root apoplastic fractions (A, B) and whole roots (C, D) of 1-week-old *ichg* mutants and WTs (n = 4). (A, C) The individual black dots indicate raw data points. The outliers were identified using the 1.5 × interquartile range rule. (B, D) The sum of daidzein, genistein, their glucoside, and malonylglucoside contents. Bar graphs show the mean of the biological replicates and circles indicate raw data points for each replicate. The Student’s two-tailed t-test was used for statistical analysis between the *ichg* mutant and WT allele from the missense and nonsense mutants, respectively (*p* < 0.05). AF, apoplastic fraction; DW, dry weight; ICHG, isoflavone conjugate-hydrolyzing β-glucosidase; WT, wild-type.

As ICHG defects affected isoflavone contents in the root apoplast, we investigated the changes of isoflavones in the root exudate of *ichg* mutants in the same growth stage. In the hydroponic medium of *ichg-1* and *ichg-2*, the contents of isoflavone glycosides significantly increased, while aglycone contents did not show a significant change from the WTs (Fig. 4A). The total isoflavone contents in the medium of *ichg* mutants were higher than in WTs due to the increase of isoflavone glycosides (*ichg-1*, *p* = 0.133; *ichg-2*, *p* < 0.001) (Fig. 4B). The total isoflavone contents in roots were comparable between the mutants and WTs, while isoflavone aglycone contents showed decreasing trends in the mutants (daidzein: *ichg-1*, *p* = 0.208; *ichg-2*, *p* = 0.013; genistein: *ichg-1*, *p* = 0.161; *ichg-2*, *p* = 0.006) (Fig. 4C, D). Together, these results indicate that ICHG involves the accumulation of isoflavones in the soybean root apoplast and their release from roots.

**Figure 4.**
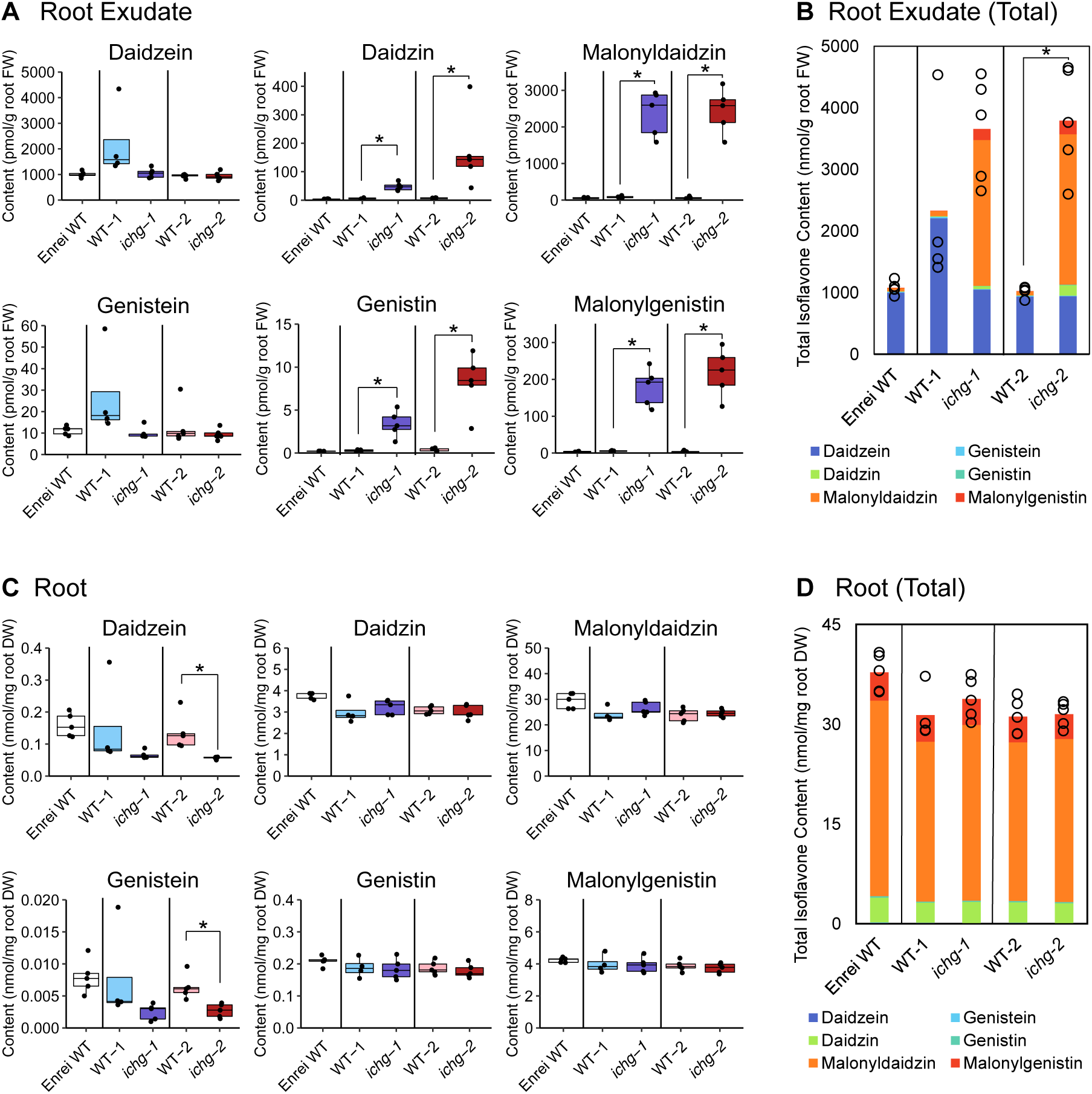
Individual and total contents of the major isoflavones in the root exudates (A, B) and roots (C, D) of 1-week-old *ichg* mutants and WTs after cultured hydroponically for 24 hours (WT-1, n = 4; others, n = 5). (A, C) The individual black dots indicate raw data points. The outliers were identified using the 1.5 × interquartile range rule. (B, D) The sum of daidzein, genistein, their glucoside, and malonylglucoside contents. Bar graphs show the mean of the biological replicates and circles indicate raw data points for each replicate. The Student’s two-tailed t-test was used for statistical analysis between the *ichg* mutant and WT allele from the missense and nonsense mutants, respectively (*p* < 0.05). DW, dry weight; FW, fresh weight; ICHG, isoflavone conjugate-hydrolyzing β-glucosidase; WT, wild-type.

### Transcriptome analysis of field-grown ichg mutants

Since the genome of the *ichg* mutants and WTs derived from the EMS mutant library has mutations in other genes than *ichg*, we assessed the impact of the mutations on the transcriptomes of WT-1, WT-2, *ichg-1*, and *ichg-2*. We conducted RNA-seq analysis using total RNA collected from 7-week-old field-grown soybean leaves and roots. The *ICHG* gene expression levels in roots were approximately more than 10-fold higher than those in leaves and they showed no significant differences between the *ichg-1* and WT-1 in both organs (Fig. 5A); whereas, in *ichg-2* the expression level of *ICHG* was significantly lower than in WT-2 (leaf, *p* = 0.002; root, *p* = 0.005) (Fig. 5A). Principal component analysis (PCA) of the transcriptomic data showed no statistically distinctive characteristics in any of the genotypes (Fig. 5B, Supplementary Table S1). The numbers of differentially expressed genes (DEGs) between WT-1 and *ichg-1*, and WT-2 and *ichg-2* are provided in Supplementary Table S2. No common genes were found in the DEGs when comparing WT-1 to *ichg-1* and WT-2 to *ichg-2* (Supplementary Dataset 1, 2). As for the isoflavone biosynthetic genes, only the expression of the *isoflavone reductase homolog 2* (*Glyma.04G012300*) was found to be up-regulated in *ichg-1* leaves (Supplementary Dataset 1) while its gene expression did not increase in *ichg-2*. Taken together, these results demonstrate that the loss of ICHG activity and other possible mutations in the genome of the mutants did not largely affect the soybean transcriptome or isoflavone metabolism.

**Figure 5.**
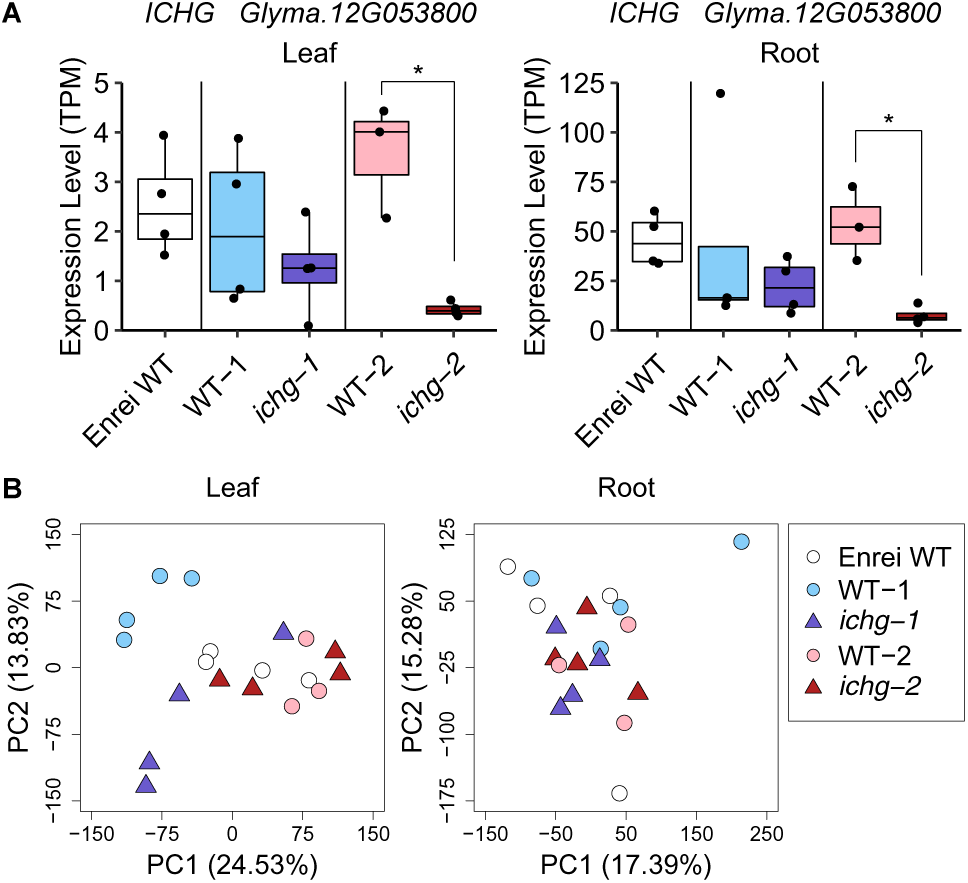
Transcriptome comparison of field-grown *ichg* mutants and WTs (WT-2, n = 3; others, n = 4). (A) mRNA levels of *ICHG* in the leaves and roots. The individual black dots indicate raw data points. The outliers were identified using the 1.5 × interquartile range rule. Student’s two-tailed t-test was used for statistical analysis between the *ichg* mutant and WT allele from the missense and nonsense mutants, respectively (*p* < 0.05). (B) Scatter plot of the first two PCs in the principal component analysis of the leaf and root transcriptomes. Circles and triangles indicate the *ICHG* WTs and *ichg* mutants, respectively. ICHG, isoflavone conjugate-hydrolyzing β-glucosidase; PC, principal component; TPM, transcripts per million; WT, wild-type.

### Isoflavone contents in field-grown ichg mutants

To examine how ICHG defects affect the isoflavone contents and composition in field-grown soybeans, we extracted isoflavone aglycones and glycosides from the leaves, roots, and rhizosphere soil. In the leaves, daidzein and genistein were not detected (Fig. 6A), and their glucosides and malonylglucosides contents showed no significant differences between the WT-1 and *ichg-1* or WT-2 and *ichg-2* (Fig. 6A, B). In roots, the daidzein and genistein contents in *ichg-1* and *ichg-2* were remarkably lower than those in WT-1 and WT-2, respectively (daidzein: *ichg-1*, *p* = 0.025; *ichg-2*, *p* = 0.024; genistein: *ichg-1*, *p* = 0.009; *ichg-2*, *p* = 0.005) (Fig. 6C). The isoflavone glycosides contents did not significantly differ between the *ichg* mutants and WTs (Fig. 6C). The total isoflavone content in the roots of both *ichg* mutants was slightly higher than that of their corresponding WTs (Fig. 6D), but the results were not statistically significant.

**Figure 6.**
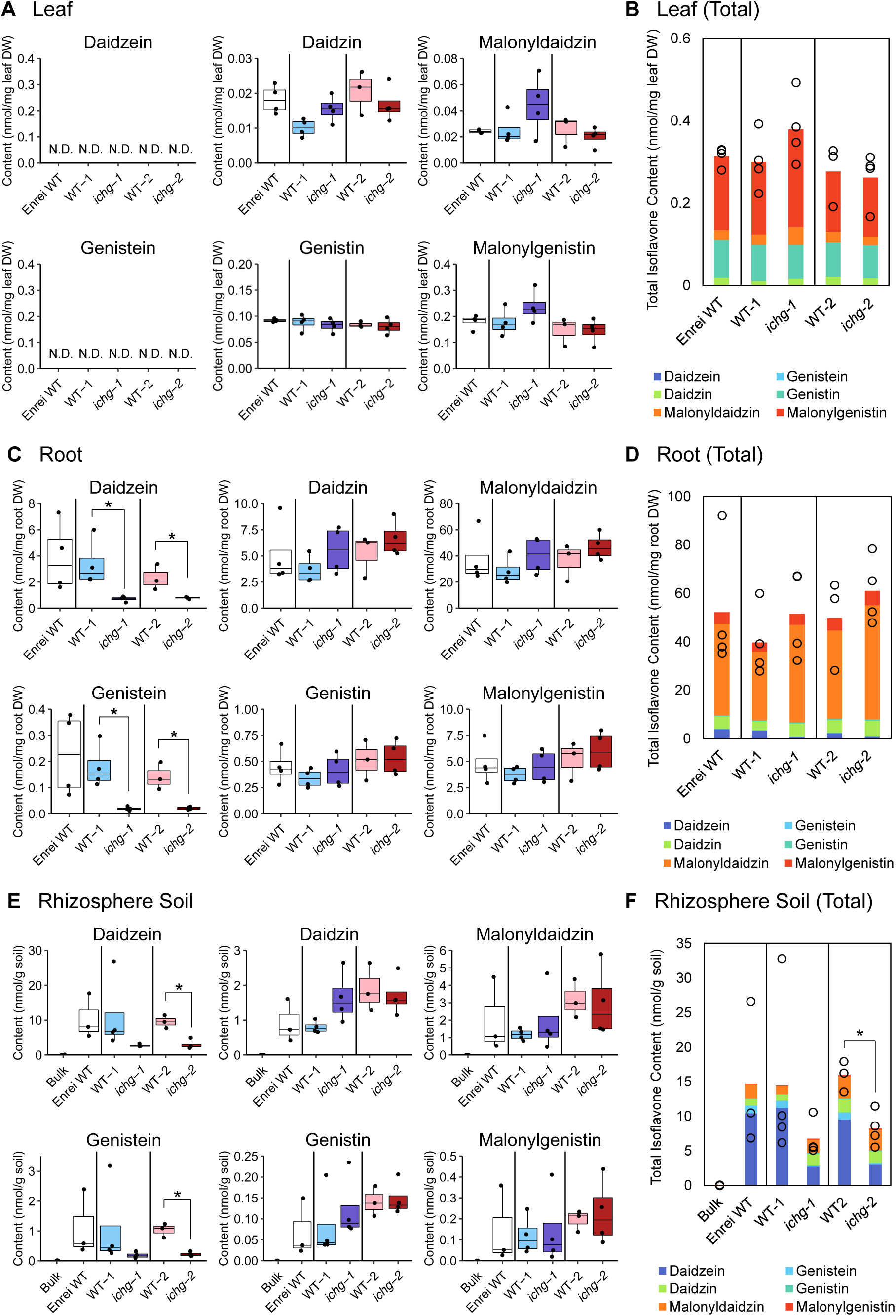
Individual and total contents of the major isoflavones in the leaves (A, B), roots (C, D), and rhizosphere soil (E, F) collected from field-grown *ichg* mutants and WTs (WT-2, n = 3; others, n = 4). Rhizosphere samples of Enrei WT lacked one replicate due to a technical error. (A, C, E) The individual black dots indicate raw data points. The outliers were identified using the 1.5 × interquartile range rule. (B, D, F) The sum of daidzein, genistein, their glucoside, and malonylglucoside contents. Bar graphs show the mean of biological replicates and the circles indicate raw data points for each replicate. Student’s two-tailed t-test was used for statistical analysis between the *ichg* mutant and WT allele from missense and nonsense mutants, respectively (*p* < 0.05). DW, dry weight; ICHG, isoflavone conjugate-hydrolyzing β-glucosidase; N.D., not detected; WT, wild-type.

In the rhizosphere soil, the amount of daidzein and genistein in *ichg-2* was significantly lower than in WT-2 (daidzein, *p* = 0.003; genistein, *p* = 0.001) (Fig. 6E). The isoflavone aglycone content in *ichg-1* was also decreased when compared with WT-1 (Fig. 6E). As the contents of the isoflavone glycosides were comparable between the mutants and WTs, the total amount of isoflavones in the rhizosphere soil of *ichg-1* and *ichg-2* was reduced to approximately half of that in the rhizosphere soil of the WT plants (Fig. 6E, F). There were only trace amounts of the isoflavones for both aglycones and glycosides detected in the bulk soil (Fig. 6E, F). These results suggest that ICHG elevates the accumulation of isoflavones in the soybean rhizosphere.

### Endosphere and rhizosphere bacterial communities of field-grown ichg mutants

Isoflavones enrich *Comamonadaceae* in the soil microbiome and alter the bacterial communities to resemble those of the soybean rhizosphere (Okutani et al., 2020). To evaluate the effects of the change of isoflavone contents in the apoplast and rhizosphere on the microbiome, we analyzed both the endosphere and rhizosphere bacterial communities using amplicon sequencing of the V4 region of the 16S rRNA. Rarefaction curves showed a similar number of the observed sequence variants in *ichg* mutants and their corresponding WTs (Supplementary Fig. S1A). The principal coordinate analysis (PCoA) of the weighted and unweighted UniFrac distance toward all samples exhibited clear distinctions between the endosphere, rhizosphere, and bulk soil bacterial communities (Permutational multivariate analysis of variance (PERMANOVA), *q* = 0.001) (Supplementary Fig. S1B, Table S3). As for the endosphere bacterial communities, the PCoA of the weighted and unweighted UniFrac distance showed no clear distinctions among the genotypes examined (Fig. 7A, Supplementary Table S4). Similarly, no significant differences were observed among the rhizosphere bacterial communities of the examined genotypes (Fig. 7B, Supplementary Table S5). The relative abundance of *Comamonadaceae* in the *ichg-2* endosphere microbiome was lower than in WT-2 (*p* = 0.013) but there was no significant difference between WT-1 and *ichg-1* (Supplementary Fig. S1C). In the rhizosphere, the relative abundance of *Comamonadaceae* did not differ among the genotypes (Supplementary Fig. S1C). The numbers of enriched and depleted bacterial families between WT-1 and *ichg-1*, and WT-2 and *ichg-2* are provided in Supplementary Table S6. No common family was found when comparing *ichg-1* to WT-1 and *ichg-2* to WT-2 except for *Yersiniaceae*, which was enriched in the endosphere of both *ichg-1* and *ichg-2* (Supplementary Figure S1D, Dataset 3, 4). The relative abundance of *Yersiniaceae* negatively correlated with daidzein contents in the root and rhizosphere soil (Spearman’s rank correlation coefficient: root, -0.7 (*p* < 0.01); rhizosphere soil, -0.67 (*p* < 0.01)). These results suggest that ICHG is not crucial for the assembly of the soybean rhizosphere bacterial communities, however, partly affects endosphere bacterial communities.

**Figure 7.**
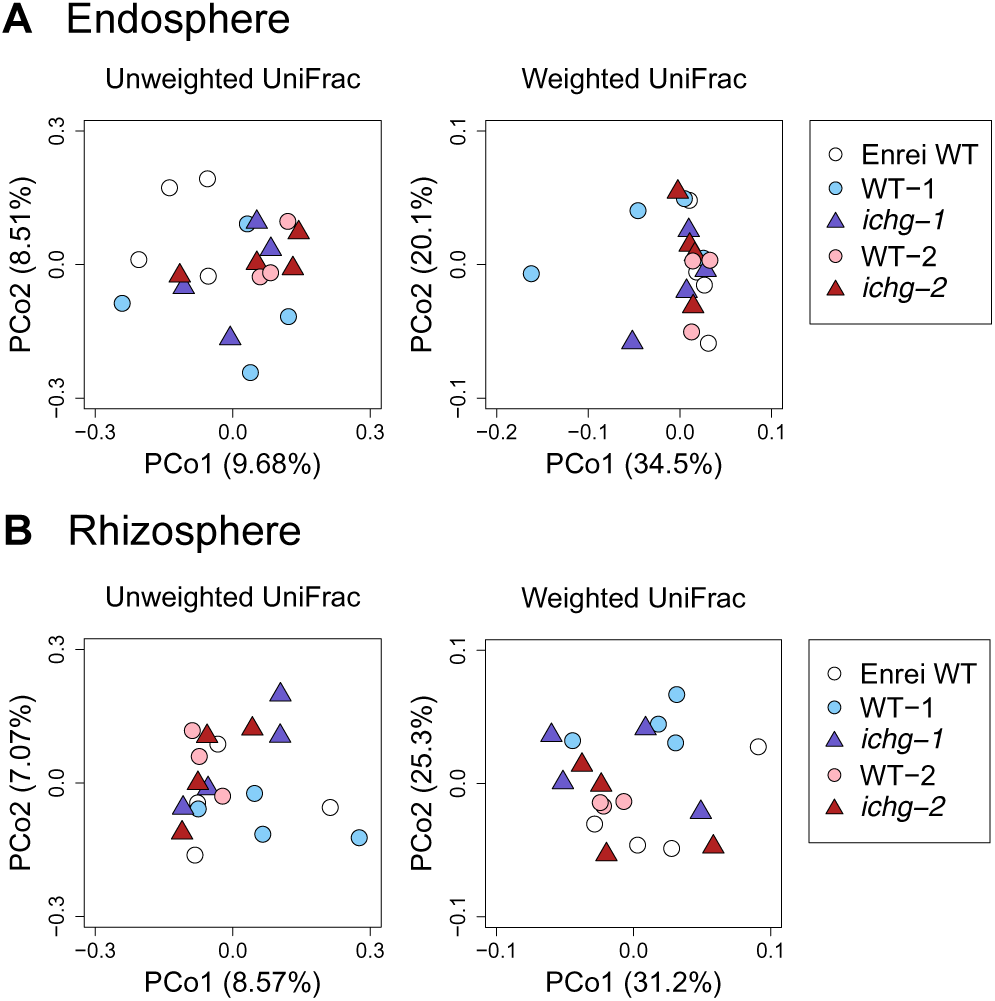
Unweighted and weighted UniFrac-based PCoA of the endosphere (A) and rhizosphere (B) bacterial communities from field-grown *ichg* mutants and WTs (WT-2, n = 3; others, n = 4). Circles and triangles indicate *ICHG* WTs and mutants, respectively. ICHG, isoflavone conjugate-hydrolyzing β-glucosidase; PCoA, principal coordinate analysis; WT, wild-type.

### Root transcriptome, isoflavone contents, and bacterial communities of ichg mutants under the nitrogen-deficient condition

Under a nitrogen-limited environment, isoflavone biosynthetic genes are up-regulated in legume roots and more isoflavone aglycones are secreted from roots (Coronado et al., 1995; Sugiyama et al., 2016). To investigate the roles of ICHG when isoflavone production and secretion are highly active, we grew *ichg* mutants both under nitrogen-sufficient (N+) and nitrogen-deficient (N-) conditions, sampled their roots at V3 stage when ICHG expression is high (Sugiyama et al., 2016). The results of PCA on the RNA-seq data showed two-separated clusters of N+ and N-samples (PERMANOVA, *q* = 0.005) (Fig. 8A, Supplementary Table S7). The DEGs comparing N+ and N-conditions are listed in Supplementary Dataset 5. Under N-condition, genes involved in isoflavone biosynthesis were up-regulated as observed in the previous studies (Chu et al., 2017; Nezamivand-Chegini et al., 2022; Sun et al., 2021) (Supplementary Figure S2, Dataset 5). The expression levels of *ICHG* in N-conditions did not increase but showed a declining trend in Enrei WT (*p* = 0.101) and *ichg-1* (*p* = 0.047) (Fig. 8B). In *ichg-2*, *ICHG* expression levels were significantly lower than WT-2 (N+, *p* = 0.004; N-, *p* = 0.023) (Fig. 8B), which were the same results with field-grown soybean leaves and roots (Fig. 5A). The contents of daidzein and genistein in N-roots showed upward trends from N+ roots in both *ichg* mutants and WTs (Fig. 8C). Comparing *ichg* mutants with WTs, the difference in the isoflavone aglycone contents was more pronounced in N-than N+ roots (Fig. 8C). As for isoflavone glycosides, no significant differences were observed except for the considerable increase in the malonyldaidzin contents in Enrei WT under N-conditions (Tukey’s HSD test*, p* = 0.018) (Fig. 8C). These results suggest that ICHG is not directly involved in the increase of the isoflavone contents in soybean roots under N-condition, however, the effect of the *ichg* mutation on the isoflavone aglycone contents becomes more apparent under N-condition than N+ condition.

**Figure 8.**
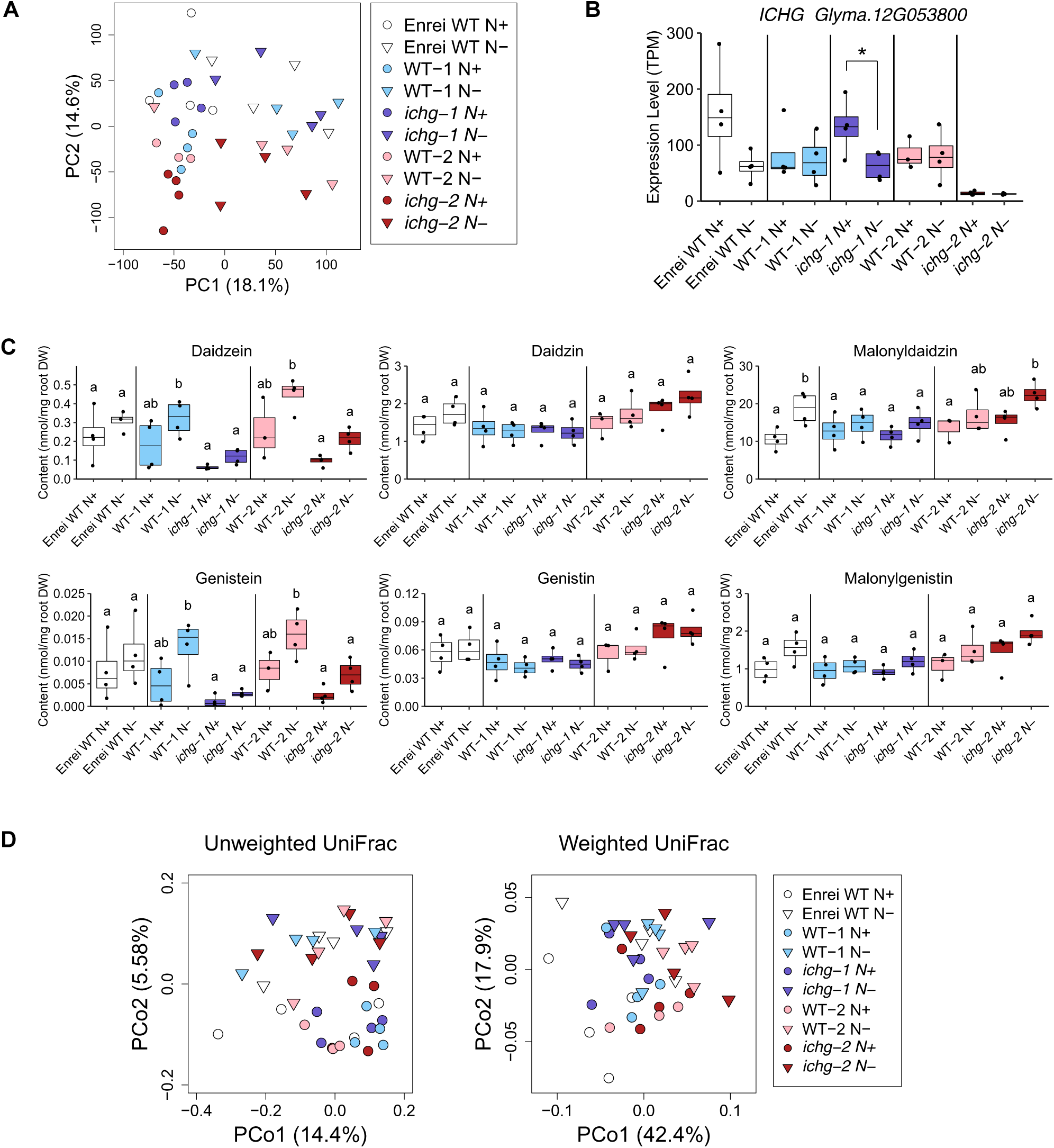
Analysis of *ichg* mutants and WTs cultivated under nitrogen-sufficient (N+) and -deficient (N-) conditions (WT-2 N+, n = 3; others, n = 4). (A) Scatter plot of the first two PCs in the principal component analysis of the transcriptome of *ichg* mutants and WTs. Circles and inverted triangles indicate samples cultivated under N+ and N-, respectively. (B) mRNA levels of *ICHG* in roots. The individual black dots indicate raw data points. The outliers were identified using the 1.5 × interquartile range rule. Student’s two-tailed t-test was used for statistical analysis between N+ and N-conditions in the same genotypes (*p* < 0.05). (C) Contents of major isoflavones in roots. The individual black dots indicate raw data points. The outliers were identified using the 1.5 × interquartile range rule. Tukey’s HSD test was used for statistical analysis among Enrei WT, the *ichg* mutant and WT allele from missense and nonsense mutants, respectively (*p* < 0.05). (D) Unweighted and weighted UniFrac-based PCoA of the endosphere bacterial communities. Circles and inverted triangles indicate samples cultivated under N+ and N-, respectively. DW, dry weight; ICHG, isoflavone conjugate-hydrolyzing β-glucosidase; PC, principal component; PCoA, principal coordinate analysis; TPM, transcripts per million; WT, wild-type.

In endosphere bacterial communities, the number of observed sequence variants did not differ among genotypes and/or nitrogen conditions (Supplementary Fig S3A). Unweighted and weighted UniFrac-based PCoA results showed a clear distinction between root bacterial communities under N+ and N-conditions (PERMANOVA, *q* = 0.001) (Fig. 8D, Table S8) and 13 genera were enriched and 26 genera were depleted under N-conditions, regardless of the ICHG defects (Supplementary Dataset 6). Relative abundance of an unannotated genus of *Comamonadaceae* and *Bradyrhizobium* showed upward trends in all examined genotypes under N-conditions (Supplementary Fig. S3B, Dataset 6). No common genus was found when comparing *ichg* mutants and WTs under both N+ and N-conditions (Supplementary Dataset 7, 8). These results suggest that ICHG is not pivotal for the assembly of the soybean root bacterial communities under both N+ and N-conditions.

### Effect of the ichg mutation on nodule formation

We analyzed whether the decrease in isoflavone aglycone contents in the root and rhizosphere affects rhizobial nodulation. Both *ichg* mutants and WTs were grown with *B. diazoefficiens* USDA 110 under N-conditions for 4 weeks. Shoot fresh weights did not show any statistically significant differences among WT-1, WT-2, *ichg-1,* and *ichg-2* (Fig. 9A). The root dry weight of *ichg-1* was higher than that of WT-1 (*p* = 0.029) (Fig. 9B). The nodule number of *ichg-2* was higher than that of WT-2 (*p* = 0.043), but there was no significant difference between WT-1 and *ichg-1* (Fig. 9C). Nodule weights were comparable across WT-1, WT-2, *ichg-1*, and *ichg-2* (Fig. 9D). The morphology of infection threads also appeared to be similar among all genotypes (Supplementary Fig. S4). Together, these results suggest that ICHG is not an essential component of the nodulation process.

**Figure 9.**
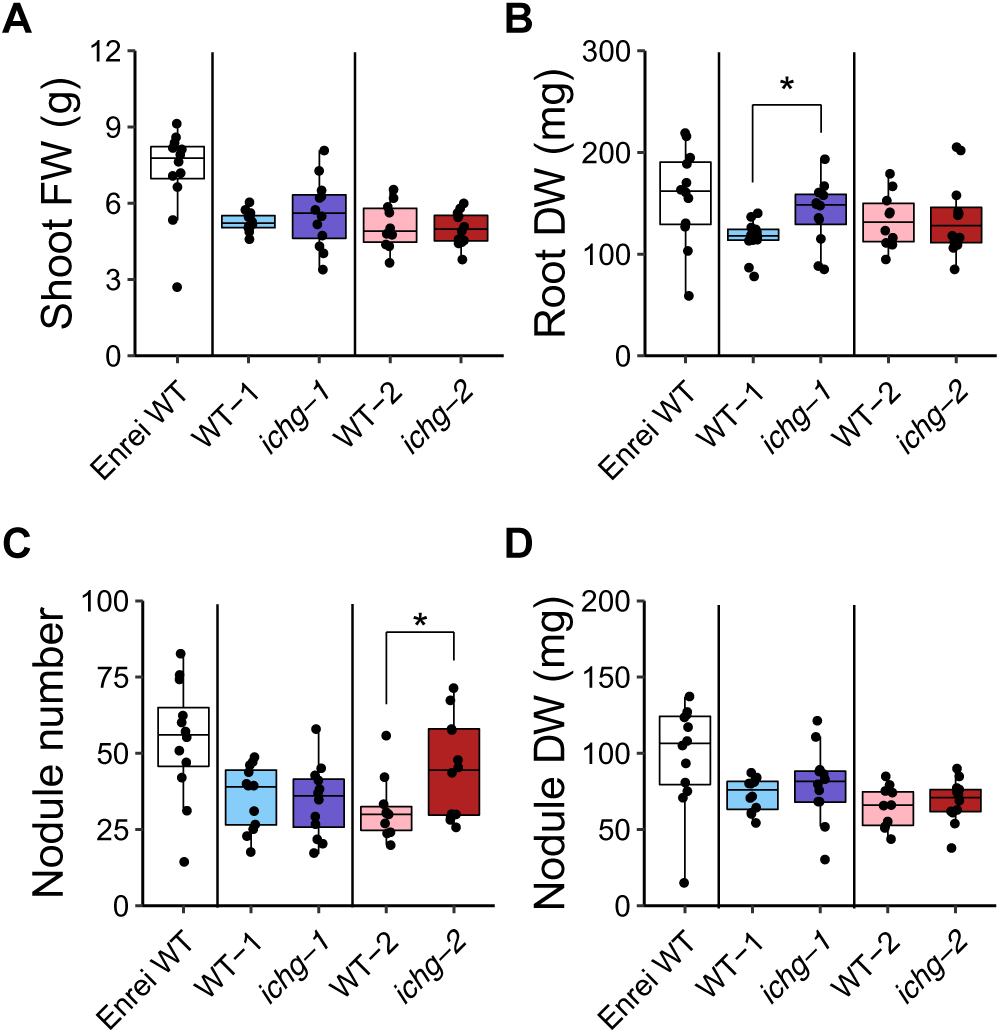
Nodulation assessment of *ichg* mutants and WTs (WT-2, n = 10; others, n = 12). (A) Fresh weight of the shoots. (B) Dry weight of the roots. (C, D) Nodule number (C) and dry weight (D). The individual black dots indicate raw data points. The outliers were identified using the 1.5 × interquartile range rule. Student’s two-tailed t-test was used for statistical analysis between the *ichg* mutant and WT allele from missense and nonsense mutants, respectively (*p* < 0.05). DW, dry weight; FW, fresh weight; ICHG, isoflavone conjugate-hydrolyzing β-glucosidase; WT, wild-type.

## Discussion

We obtained *ichg* missense and nonsense mutants by screening a soybean high-density mutant library. The *ichg-2* mutant harbored a nonsense mutation (Gln232stop) and the *ichg-1* mutant harbored a missense mutation (Glu420Lys), which resulted in the complete loss of ICHG activity, as this glutamic acid is one of two highly conserved glutamic acids among the glycosyl hydrolase family 1 β-glucosidases and a proposed catalytic residue (Barrett et al., 1995). Although there was no remarkable change in the transcriptomes of the *ichg* mutants both in the field and under the nitrogen-deficient conditions, isoflavone aglycone contents were decreased in the root, apoplastic fraction, and rhizosphere soil (Fig. 10). The expression of isoflavone biosynthetic genes and candidate transporter genes were comparable between *ichg* mutants and WTs and were up-regulated under nitrogen-deficient conditions in both *ichg* mutants and WTs (Supplementary Fig. S2, S5) (Matsuda et al., 2020), indicating that there are little feedback or feedforward regulations on isoflavone metabolism derived from the effect of *ichg* mutation.

**Figure 10.**
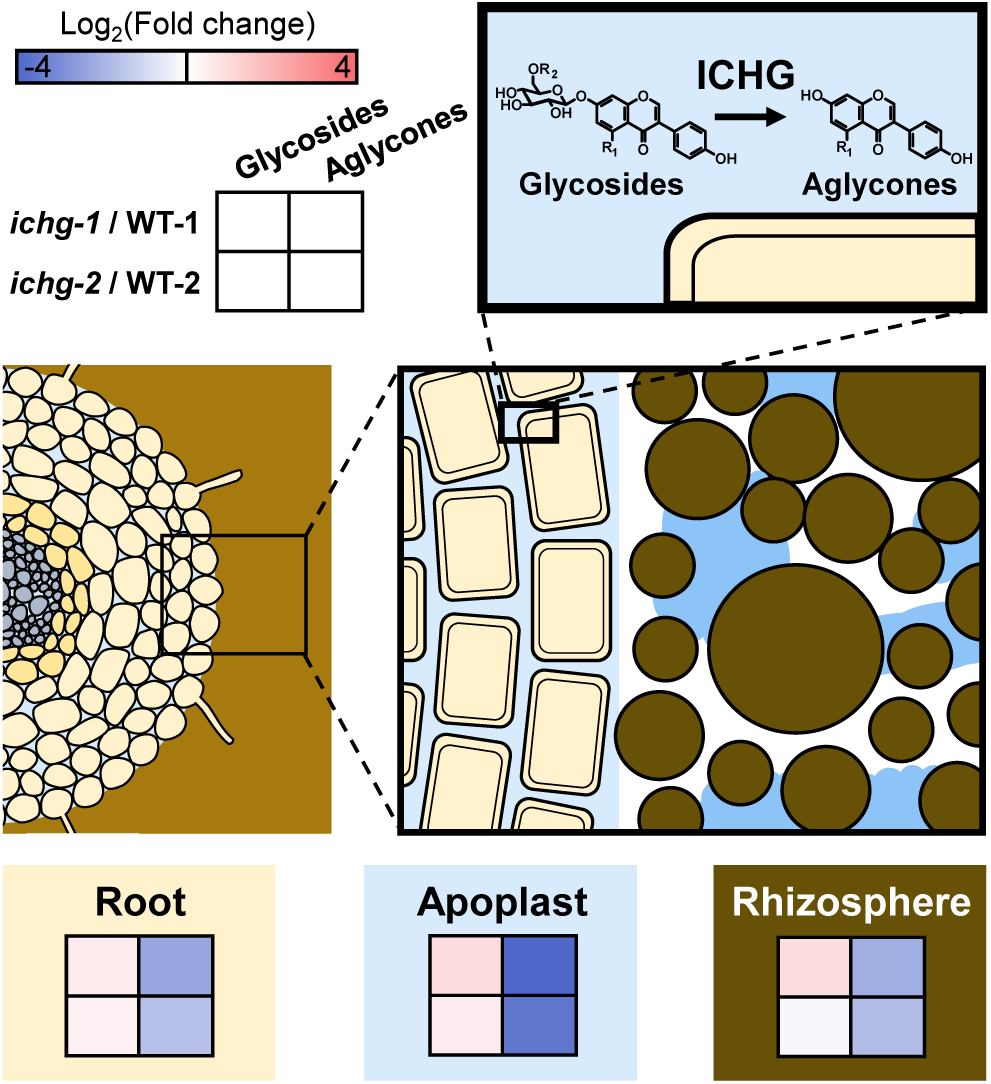
Heatmaps showing the fold change of isoflavone glycoside and aglycone contents in roots (yellow box), the apoplastic fraction (light blue box), and rhizosphere soil (brown box) of *ichg* mutants compared with those of WTs. The color of each cell corresponds to the log_2_(fold change) of the mean of isoflavone aglycone (the sum of daidzein (R_1_ = H) and genistein (R_1_ = OH)) or glycoside (the sum of their glucosides (R_2_ = H) and malonylglucosides (R_2_ = COCH_2_COOH)) contents in *ichg-1* compared to WT-1, or *ichg-2* compared to WT-2. Red and blue colors indicate the increase and decrease in *ichg* mutants against the WTs, respectively. ICHG, isoflavone conjugate-hydrolyzing β-glucosidase; WT, wild-type.

Despite a reduction in the aglycone content of the roots and rhizosphere of the *ichg* mutants, there were no remarkable common characteristics in the endosphere or rhizosphere bacterial communities or a specific nodulation phenotype. This was probably because the changes in the aglycone contents of the endosphere and rhizosphere were not drastic enough to affect the bacterial communities in the environment when other metabolites were present. For nodulation, isoflavone aglycone daidzein and genistein have been regarded as the major components of soybean root extracts responsible for inducing *nod* genes in *B. diazoefficiens* (Kosslak et al., 1987). A recent study by Ahmad et al. (2021) suggested that isoflavone malonylglucosides also induce Nod factor production based on the analysis of over-expression and knockdown mutants of an isoflavone malonyltransferase, GmMaT2. Our analysis using isoflavone standard chemicals showed the slight induction of *nodD1* gene, but not *nodA* nor *nodY* genes, by malonyldaidzin (Supplementary Fig. S6), which is consistent with the previous study showing that isoflavone glycosides can induce *B. diazoefficiens* (*B. japonicum*) *nodD1* gene but not *nodYABCSUIJ* operon (Smit et al., 1992). These results suggest that the changes in the contents of isoflavone glycosides and aglycones in the *ichg* mutants do not greatly affect the efficacy of *nod* gene induction.

Isoflavone aglycones are known to be secreted from soybean roots and function in the rhizosphere but their underlying secretion mechanisms are poorly understood (Hassan and Mathesius, 2012; Sugiyama, 2021). While plasma membrane-localized ABC and MATE transporters mediate the export of isoflavone aglycones (Biała-Leonhard et al., 2021; Sugiyama et al., 2007), there is currently no evidence to explain how apoplast-localized β-glucosidase ICHG is involved in the isoflavone supply to the rhizosphere. In general, glycosides are unstable in the soil, and those secreted from the roots are hydrolyzed to form aglycones within a few hours by soil bacterial enzymes, which has led to the assumption that ICHG has a minor role in the accumulation of isoflavones in the rhizosphere (Kong et al., 2007; Sugiyama et al., 2017; Weir et al., 2010). However, we showed that the total isoflavone content in the rhizosphere soil of *ichg* mutants was significantly decreased (Fig. 10), suggesting that the ICHG-mediated hydrolysis of isoflavone glycosides into aglycones is pivotal for the accumulation of isoflavones in the rhizosphere. While isoflavone glycoside contents in the hydroponic medium of 1-week-old soybeans were significantly increased in *ichg* mutants, those in the rhizosphere soil of 7-week-old field-grown *ichg* mutants were not increased. The increase of isoflavone glycosides in a hydroponic medium may be due to the differences in the developmental stage and/or growth conditions.

PSMs often exist in vacuoles as their glycosylated forms and the rhizosphere as their partially or fully de-glycosylated forms (Frey et al., 2009; Grubb and Abel, 2006; Neal et al., 2012; Strehmel et al., 2014; Stringlis et al., 2018; Tsuno et al., 2018). In *Arabidopsis*, coumarin glucoside scopolin is accumulated in roots and its aglycone, scopoletin, is secreted from roots under iron-limited conditions, in which this coumarin modulates the rhizosphere microbiota for systemic resistance and iron uptake (Stringlis et al., 2018). An *Arabidopsis* β-glucosidase, BGLU42, predicted to be localized in plastids, hydrolyzes scopolin to form scopoletin (Hooper et al., 2017). BGLU42 is required for the release of scopoletin into the rhizosphere, as *bglu42* mutants were found to secrete fewer scopoletin and instead accumulated larger amounts of scopolin in the roots (Stringlis et al., 2018; Zamioudis et al., 2014). Benzoxazinoids and glucosinolates are also found mostly in their deglycosylated forms in the rhizosphere (Neal et al., 2012; Strehmel et al., 2014), but whether de-glycosylating enzymes are involved in their secretion, and which specific enzymes may be involved, requires further investigation. The present study sheds light on the previously unrecognized role of apoplast-localized β-glucosidases in relation to the accumulation of PSMs in the rhizosphere. Since multiple secretory routes bring PSMs to the rhizosphere, it is important to elucidate how each route functions coordinately under various stress and developmental conditions. Further studies on β-glucosidases, other de-glycosylating enzymes, and other proteins related to PSM secretions will enable us to better understand the dynamics and functions of PSMs in the rhizosphere.

## Materials and methods

### Chemicals and plant materials

Malonyldaidzin and malonylgenistin were purchased from Nagara Science. All other chemicals were purchased from Wako Pure Chemical Industries Ltd. or Nacalai Tesque Inc., unless otherwise stated. Seeds of WT soybean cv Enrei (Enrei WT) were purchased from Tsurushin Syubyo, Matsumoto, Japan.

### Screening ichg mutants from an EMS mutant library

The *ichg* mutants were obtained from a high-density mutant library of the cultivar Enrei according to a previously described amplicon sequencing method (Tsuda et al., 2015). This library consisted of DNA and seeds from 1,536 EMS-induced mutant plants. Amplicons of the *ICHG* gene (*Glyma.12G053800* for Gmax_275 reference) which were approximately 6,700 bp, were amplified from 384 bulk DNA samples, each consisting of material from four mutant plants. PCR reaction mixtures (10 μL) contained 0.2 μL of template DNA from each of the 384 DNA pools, 2 μL of 5× PrimeSTAR GXL Buffer (Takara Bio, Kusatsu, Japan), 1.0 μL of PrimeSTAR GXL DNA Polymerase (1.25 U/μL), 0.8 μL of 2.5 mM dNTPs, and 0.1 μL of each of the 20 μM forward and reverse primers [ICHG-AS_F (5’-aatttggaatccgtgagtttcttgtga-3’) and ICHG-AS_R (5’-taataattcccgtcttgctttgtgctt-3’)]. PCR was performed on a GeneAmp PCR System 9700 (Applied Biosystems, Foster City, CA, USA) with the following program: initial denaturation for 5 s at 98℃; 30 cycles of denaturation for 10 s at 98℃, annealing for 15 s at 68℃, and extension for 7 min 35 s at 68℃; and a final extension for 30 s at 68℃. Four PCR samples were mixed to prepare 96 PCR amplicon pools. The dual index sequencing library was prepared using a Nextera™ XT DNA Sample Preparation Kit and Nextera™ XT Index Kit (Illumina, San Diego, CA, USA), both according to the manufacturers’ instructions. Paired-end sequencing data was obtained on a MiSeq™ platform using a MiSeq™ v2 500-cycle kit (Illumina) with the default parameters. Read mapping and variant detection of the 96 PCR amplicon pools were conducted using CLC Genomics Workbench software (CLC Bio, Aarhus, Denmark) with the workflow and batch processing tools, according to the parameter settings described by Tsuda et al. (2015). PCR amplicon pools that contained missense (Glu420Lys hereafter *ichg-1*) or nonsense (Gln232stop hereafter *ichg-2*) mutations in the *ICHG* gene were identified using the variant detection procedure (Tsuda et al., 2015). The plants in which the mutations occurred were identified by directly sequencing each of the 16 original DNA samples from the PCR amplicon pool. Direct sequencing was conducted using primer pairs (5’-attataatgcaggccgcttcagtttg-3’ and 5’-agagcttcttccacggaaagggttg-3’ for *ichg-1*; and 5’-ttgatatgattacaagttgttgagctttg-3’ and 5’-cttagtcttgtacacatgaacagcagc-3’ for *ichg-2*), a BigDye Terminator v3.1 Cycle Sequencing Kit, and an ABI 3730xl genetic analyzer (Thermo Fisher Scientific, Waltham, MA, USA) according to the manufacturers’ instructions. The mutants *ichg-1* and *ichg-2* were backcrossed with the WT cultivar Enrei, and the self-pollinated F_2_ progenies were used in the following experiment. The genotype of the F_2_ progenies was determined by sequencing the amplicons obtained using a primer pair (5’-aggtctaagggtacatttgtag-3’ and 5’-tcttccacggaaagggttg-3’ for *ichg-1*; 5’-actaaaaatgttcgaaattcg-3’ and 5’-caactcaaagttgtgtgagaaaca-3’ for *ichg-2*). The seeds from the F_2_ plants that were homozygous for the mutants or wild alleles (*ichg-1* and WT-1; *ichg-2* and WT-2) were used in the following experiments.

### Extraction of apoplastic crude proteins

Soybean seeds (Enrei WT, WT-1, *ichg-1*, WT-2, and *ichg-2*) were sterilized with chlorine gas for 3 hours and germinated in plant boxes (AGC Techno Glass, Haibara, Japan) filled with autoclaved vermiculite and water (all sterilization and germination methods were the same hereafter unless otherwise specified). The 1-week-old seedlings were rinsed and transferred to a hydroponic culture system as previously described (Matsuda et al., 2020). Approximately 3 g of fresh roots from three individual 34-day-old soybean plants of each genotype and 2.5 g of polyvinylpolypyrrolidone were homogenized in a mortar with 15 mL of the homogenizing buffer [100 mM phosphate buffer (pH 7.5), 1 mM dithiothreitol (DTT), and 1 mM phenylmethylsulfonyl fluoride (PMSF)]. After filtering with Miracloth (Merck, Darmstadt, Germany), the pellet was suspended with 15 mL of a buffer for apoplastic protein extraction [100 mM phosphate buffer (pH 7.5), 1 mM DTT, 1 mM PMSF, 2 M NaCl] and incubated on ice for 1 hour with stirring every 15 min. The suspension was filtered with Miracloth (Merck) and the supernatant was centrifuged (5,800 *g*, 4℃, 10 min). Then, 3 mL of the centrifuged supernatant was applied to a PD-10 column filled with Sephadex G-25 resin (GE Healthcare, Chicago, IL, USA) equilibrated with the homogenizing buffer. Then 3 mL of the homogenizing buffer was applied to the column and the eluate collected and stored at −80℃ until further use.

### In vitro ICHG assay

Extracted apoplastic crude proteins (10 µg) were mixed with the homogenizing buffer to a volume of 95 µL on ice. After incubating the mixture at 30℃ for 3 min, 5 µL of 1 mM malonyldaidzin in methanol was added with gentle tapping and centrifugation. The solution was then incubated at 30℃ for 20 min, and the reaction was stopped with 200 µL of 1 % (v /v) formic acid in methanol, followed by centrifugation (20,400 *g*, 4℃, 15 min). The supernatant was filtered with Minisart RC4 (pore size 0.20 µm, diameter 4 mm, Sartorius stedim biotech, Göttingen, Germany) and injected into a high-performance liquid chromatography system (LC-10AD, Shimadzu, Kyoto, Japan) with TSKgel ODS-80TM (4.6 × 250 mm, 5 µm, TOSOH Corporation, Tokyo, Japan) to quantitate the daidzein. The LC mobile phase consisted of (A) water and (B) acetonitrile, which both contained 0.3 % (v /v) formic acid, and was eluted isocratically at 36 % of (B) solution for 20 min. The flow rate was 0.8 mL/min and daidzein was detected at 260 nm. Experiments were performed with three technical replicates for all genotypes examined.

### Extraction of isoflavones in root apoplastic fractions and root exudates from soybean seedlings

Apoplastic fractions of the soybean roots were extracted as described previously with some modifications (Li, 2011). One-week-old soybean seedlings were harvested and gently washed twice with deionized water to remove the vermiculite. The roots were cut off, weighed, and then placed into 10 mM sodium phosphate buffer (pH 6.0) in a desiccator and vacuumed on ice. The air pressure in the desiccator was kept under –0.09 MPa for 30 min, then slowly returned to atmospheric pressure. The infiltrated roots were dabbed onto a piece of paper to dry and then placed into the plunge barrel of a 10 mL syringe. The syringe barrel was placed in a 50 mL tube and centrifuged at 3,000 *g* for 15 min at 4℃. The collected apoplastic fraction was loaded through a Sep-Pak C18 Plus Short cartridge (Waters). The cartridge was washed with 10 mL of deionized water and eluted with 1.6 mL of methanol. Eluted fractions were dried under nitrogen gas at 50℃, reconstituted in 200 µL methanol, and filtered through a 0.45 µm Minisart RC4 filter (Sartorius).

Root exudates from one-week-old soybean seedlings were collected as described previously (Sugiyama et al., 2016). After 24 hours of incubation in a nitrogen-deficient mineral nutrient medium at 24℃ under 16 h light /8 h dark cycle, the roots were cut off and stored at –80℃ for isoflavone extraction.

The extracts were injected into an LC system (ACQUITY H-Class System, Waters, Milford, MA, USA) with ACQUITY UPLC BEH C18 Column (2.1 × 50 mm, 1.7 μm, Waters). The LC mobile phase consisted of (C) water containing 0.1 % (v/v) formic acid and (D) acetonitrile. The gradient program was isocratic at 10 % D, Initial; linear at 0 %–85 % D, 0–15 min; isocratic at 100 % D, 15–16 min; and isocratic at 100 % D, 16–20.5 min. The flow rate was 0.2 mL/min. Isoflavones were detected using a tandem quadrupole MS (Xevo TQD, Waters) and the Multiple Reaction Monitoring (MRM) mode. MRM conditions for this analysis were the same as previously described (Matsuda et al., 2020).

### Cultivating and sampling field-grown soybean

Field cultivation of soybean was carried out at the Kyoto University of Advanced Science at Kameoka, Kyoto, Japan (34°99’38”N, 135°55’14”E). Surface-sterilized soybean seeds were sown in pods filled with vermiculite. Seedlings were grown at 23℃ under a 16-h-light/8-h-dark cycle for 8 days and then planted in a field on June 25, 2020. The sampling of 7-week-old soybean plants (R1 stage) was conducted on August 5, 2020. Four soybean plants for each genotype (WT, *ichg-1*, *ichg-2*, and WT-1) were sampled except for WT-2 which was collected for three plants. The upper fully expanded leaves and a few lateral roots were taken from each sample, immediately frozen in dry ice, and then transferred to –80℃ until further use. Bulk soil was collected at four different locations at least 20 cm from the plants as previously described (Sugiyama et al., 2014). Whole soybean plants were brought back to the laboratory on ice. The rhizosphere soils for isoflavone analysis were then collected from each plant using brushes, as described previously (Sugiyama et al., 2014). One replicate of Enrei WT rhizosphere samples lacked due to a technical error. Roots were then washed and sonicated in a beaker filled with 100 mL PBS buffer containing 0.02 % Silwet L-77 for 5 min each (Bulgarelli et al., 2012). The washing buffer was then centrifuged at 8,000 g for 10 min at 5℃, and the soil was collected for bacterial community analysis. The sonicated roots were rinsed with tap water and then stored at –30℃ for DNA extraction.

### Cultivating soybeans under nitrogen-sufficient and -deficient conditions

This experiment was conducted according to (Yazaki et al., 2021) with minor modifications. One-week-old soybean seedlings were transferred to pots filled with vermiculite. Approximately 4 g of soil, collected from the field at the Kyoto University of Advanced Science, Kameoka, Kyoto, Japan, was placed on the surface of vermiculite as a bacterial inoculation source. Soybean plants were cultivated in a greenhouse at 28℃ under natural light conditions at Uji campus, Kyoto University (34°90’87”N, 135°80’22”E) from June 20 to July 6, 2022, and harvested at the V3 stage. Four plants for each genotype were grown in both nitrogen-sufficient and nitrogen-deficient conditions, except for WT-2 under the nitrogen-sufficient condition which lacked one replicate due to a technical failure. After removing vermiculite from roots with tap water, nodules were removed, and some of the lateral roots were immediately cut off and frozen with liquid nitrogen for RNA and metabolite extraction. The surface of the remaining roots was washed and sonicated in a PBS buffer with 0.02 % Silwet L-77 for 5 min each and was rinsed with tap water (Bulgarelli et al., 2012). Lateral roots approximately 10 cm from the base were cut off and stored at –30℃ for DNA extraction.

### RNA-seq and transcriptome analysis

RNA extraction, sequencing, raw-read alignment, normalization, and gene annotation were performed as previously described (Matsuda et al., 2020). The DEGs between the *ICHG* WTs and mutants were determined using the DESeq2 package (Love et al., 2014) in the R environment with a false discovery rate cutoff of 0.01.

### Extraction of isoflavones from plant tissues and soil

The preparation of leaf and root extracts was performed according to (Sugiyama et al., 2016). Isoflavone extraction from the rhizosphere and bulk soil was conducted using a previously described method with some modifications (Sugiyama et al., 2017). In brief, soil samples (approximately 200 mg) were extracted in 3 × 1 mL of methanol at 50℃ (10 min each) and centrifuged at 5,000 *g* for 5 min. The combined supernatant was dried under a nitrogen stream at 50℃, re-dissolved in 100 µL methanol, and filtered through a 0.45 µm Minisart RC4 filter (Sartorius). The extracts were analyzed by LC-MS as described above.

### Bacterial community analysis using 16S rRNA gene amplicon sequencing

Bacterial community analysis was performed using 16S rRNA gene amplicon sequencing, as previously described with some minor modifications (Nakayasu et al., 2021). Briefly, the V4 region of the 16S rRNA genes was amplified using PCR with the following forward and reverse primers: 515F (5’-acactctttccctacacgacgctcttccgatct-gtgccagcmgccgcggtaa-3’) and 806R (5’-gtgactggagttcagacgtgtgctcttccgatct-ggactachvgggtwtctaat-3’), respectively, and sequenced using the MiSeq platform (Illumina). The sequence data were analyzed using QIIME2 version 2021.11 (Bolyen et al., 2019). The 21^st^ to 200^th^ base pair of paired-end sequencing reads were trimmed out and error-corrected amplicon sequence variants (ASVs) were constructed using DADA2 (Callahan et al., 2016) with the q2-dada2 plugin in QIIME2. The α- and β-diversity metrics were estimated with 50,000, 60,000, and 31,500 sequences per rhizosphere and endosphere sample of field-grown soybeans, and endosphere sample of greenhouse-grown soybeans, respectively. Taxonomic assignment of the ASVs was performed using the Naïve Bayes classifier with SILVA release 138 (Bokulich et al., 2018; Quast et al., 2013). The linear discriminant analysis (LDA) effect size (LEfSe) method (Segata et al., 2011) was performed for bacterial families or genera with more than 0.1% of the maximum relative abundance using default parameters to detect differentially abundant bacterial families with an adjusted *p*-value < 0.05 for Kruskal-Wallis and an LDA score of >2.

### Nodulation test and observation of rhizobial infection threads

*B. diazoefficiens* USDA110 (MAFF 303215) was obtained from the NARO Genebank (http://www.gene.affrc.go.jp/index_en.php). The DsRed-labeled *B. diazoefficiens* USDA 110 was created as previously described (Nguyen et al., 2020). DsRed-non-labeled and labeled *B. diazoefficiens* USDA 110 was pre-cultured in YEM medium (Estrella et al., 2004). After 7 days, 1 mL of the culture solution was centrifuged (13,000 *g*, 4℃, 2 min) and the pellet was suspended with 1 mL of 10 mM MgCl_2_. The washing process was repeated twice, and the washed suspension was then diluted to 1 × 10^7^ colony forming units (CFU)/1 mL. Ten surface-sterilized soybean seeds were then germinated in a Petri dish with 20 mL sterile water and two pieces of KimWipe (Kimberly-Clark Corp., Irving, TX, USA).

For the nodulation test, all work was staggered by one day for WT-1 and *ichg-1*, Enrei ET, and WT-2 and *ichg-2*. After 5 days, seedlings were transferred into Leonard jars made of two plant boxes (AGC TECHNO GLASS Co., Ltd.) (Ye et al., 2005) and *B. diazoefficiens* (1 × 10^7^ CFU) was inoculated onto the seedlings. The soybeans were cultivated in a greenhouse at Uji campus, Kyoto University (34°90’87”N, 135°80’22”E) from November 3–5, 2021 to December 1–3, 2021. When the bottom side of the Leonard jars became empty, 1/2 nitrogen-free medium (Yazaki et al., 2021) was poured into the jar. The room temperature was set at 23℃ with supplemental lighting in the morning and evening to create 16-h-light/8-h-dark conditions. After 28 days, 12 soybean plants for each genotype (WT, *ichg-1*, *ichg-2*, and WT-1), and 10 plants for WT-2 were sampled. The soybean shoots were cut and weighed. Their roots were gently washed with tap water and then patted dry. Nodules on the roots were picked using tweezers and counted. The leftover roots and nodules were completely dried at 50℃ and then weighed.

For the rhizobial infection thread observations, soybean seedlings were transferred to autoclaved plant boxes (AGC TECHNO GLASS Co., Ltd.) five days after germination and the DsRed-labeled *B. diazoefficiens* (1 × 10^7^ CFU) was inoculated onto the seedlings. Each plant box was covered with another sterile plant box to maintain sterility inside the box. After 7 days of incubation at 25℃ with 16-h-light/8-h-dark conditions, the roots were washed gently with deionized water. The upper part (<3 cm) of the lateral roots was cut and observed. Fluorescence images of the infection threads were captured using a confocal laser-scanning microscope (FV3000, Olympus, Tokyo, Japan) with a 40 × 0.75 numerical aperture objective. DsRed was monitored by excitation at 561 nm with a 20-mW diode laser and emission at 570– 670 nm. Three shots of different infection threads were taken for each genotype.

### Expression analysis of rhizobium nod genes

*B. diazoefficiens* USDA110 was pre-cultured in YEM medium (Estrella et al., 2004) at 28℃ with 180 rpm until OD_600_ reached 0.5∼1.0. The pre-culture solution was diluted to OD_600_ = 0.1 and dispensed at 4 mL each for four replicates, and added 4 µL of 10 mM daidzein, daidzin, and malonyldaidzin standards dissolving in dimethyl sulfoxide, and dimethyl sulfoxide as a mock treatment. After 20 hours of the *nod* gene induction at 28℃ with 180 rpm, 2 mL of the culture solution was transferred to a 2.0 mL tube and subjected to centrifugation (15,000 *g*, 4℃, 2 min). The pellet was suspended with 1 mL of TRI Reagent® (Molecular Research Center Inc., USA) by pipetting and RNA extraction was performed according to the manufacturer’s protocol. The synthesis of cDNA from 200 ng of total RNA was conducted by using ReverTra Ace^®^ qPCR RT Master Mix with gDNA Remover (Toyobo Co., Ltd., Osaka, Japan). For qPCR, 1 µL of synthesized cDNA (×1/4 dilution) was added as a template to THUNDERBIRD^®^ SYBR^®^ qPCR Mix (Toyobo Co., Ltd., Japan). The qRT-PCR reactions were performed and primer sets for amplifying *nodY*, *nodD1*, *ParA* of *B. diazoefficiens* USDA 110 were designed according to (Ahmad et al., 2021; Sugawara and Sadowsky, 2013). Relative gene expression was calculated with the Pfaffl method (Pfaffl, 2001) using *ParA* as a reference gene.

### Statistical analysis

All boxplots in this study were created in the R environment (v. 4.1.2; R Core Team, 2021) using the ggplot2 package and the outliers were identified with a default parameter (1.5 × interquartile range). The statistical analysis of plant transcriptomic data was conducted using the adonis function in the vegan R package. Since the F_2_ progenies of two independent *ichg* mutants screened from the mutant library are likely to have different genetic backgrounds, the Student’s two-tailed t-test was used for calculating statistically significant differences (\**p* < 0.05) between the missense mutant-derived the *ichg-1* and WT-1 and the nonsense mutant-derived the *ichg-2* and WT-2, respectively. Spearman’s rank correlation coefficient and its statistical test were performed in the R environment. In the nitrogen-controlled experiment, the Student’s two-tailed t-test was used for comparing N+ samples with N-samples in the same genotypes (\**p* < 0.05, \*\**p* < 0.01, \*\*\**p* < 0.001). Tukey’s HSD test was used for the analysis of isoflavone contents under N+ and N-conditions among Enrei WT, the *ichg* mutant and WT allele from missense and nonsense mutants, respectively (*p* < 0.05). The statistical analysis of bacterial communities was conducted using unweighted and weighted UniFrac-based PERMANOVA in QIIME2. Dunnett’s test was used for statistical analysis on daidzein, daidzin, and malonyldaidzin-treated samples compared to the DMSO-treated sample (\**p* < 0.05).

## Data Availability

The soybean transcriptome and 16S rRNA amplicon sequencing datasets supporting the results of this study are publicly available at the DNA Data Bank of Japan (DDBJ) (https://www.ddbj.nig.ac.jp) (DRA015007, DRA015099, DRA013654, DRA015072). The other data underlying this article will be shared on reasonable request to the corresponding author.

## Funding

This work was supported by a JSPS Research Fellowship for Young Scientists DC1 (21J23155 to H.M.); a JST-CREST grant (JPMJCR15O2 to A.J.N., JPMJCR17O2 to A.S.); JSPS KAKENHI (19H02860 to S.O., 18H02313 and 21H02329 to A.S.); the Ministry of Agriculture, Forestry, and Fisheries of Japan (Genomics-based Technology for Agricultural Improvement, IVG1005 to A.K, SFC2001 to A.S.); and a grant from the Research Institute for Sustainable Humanosphere (Mission 1), Kyoto University.

## Disclosures

Conflicts of interest: No conflicts of interest declared.

## Acknowledgments

*B. diazoefficiens* USDA110 (MAFF 303215) was provided from NARO Genebank, Japan. We thank Ms. Keiko Kanai, Ms. Yuko Kobayashi, Ms. Rie Mizuno, Ms. Kyoko Takamatsu, and Ms. Kyoko Mogami for technical assistance; Dr. Jiro Sekiya for supporting our field experiment; Dr. Toru Nakayama, Dr. Seiji Takahashi, and Dr. Toshiyuki Waki for helpful discussion; Dr. Ryosuke Munakata for critical reading of the manuscript; Ms. Haruko Hirukawa for creating the graphic of soybean. We also thank DASH/FBAS, the Research Institute for Sustainable Humanosphere, Kyoto University.

## Author contributions

H.M. and A.S. conceived and designed the research; K.Y. and A.S. supervised the experiments; A.K. screened and backcrossed *ichg* mutants; H.M., M.N., H.T., and A.S. grew and sampled soybeans for the experiments; H.M., Y.Y., E.M., and A.S. performed the protein activity assays; H.M. and A.J.N. conducted RNA-seq experiments; H.M, S.Y., and Y.A. analyzed the transcriptomic data; H.M. performed the analysis of isoflavone contents, bacterial communities, and rhizobial gene expressions; H.M. and S.O. analyzed infection threads using fluorescent-tagged rhizobia; H.M., M.N., A.K., K.Y., and A.S. wrote the article with contributions from all the authors; A.S. agrees to serve as the author responsible for contact and ensures communication.

## Notes

### Competing Interest Statement

The authors have declared no competing interest.

