## Supplemental Figures for "Apoplast-localized β-Glucosidase Elevates Isoflavone Accumulation in the Soybean Rhizosphere"

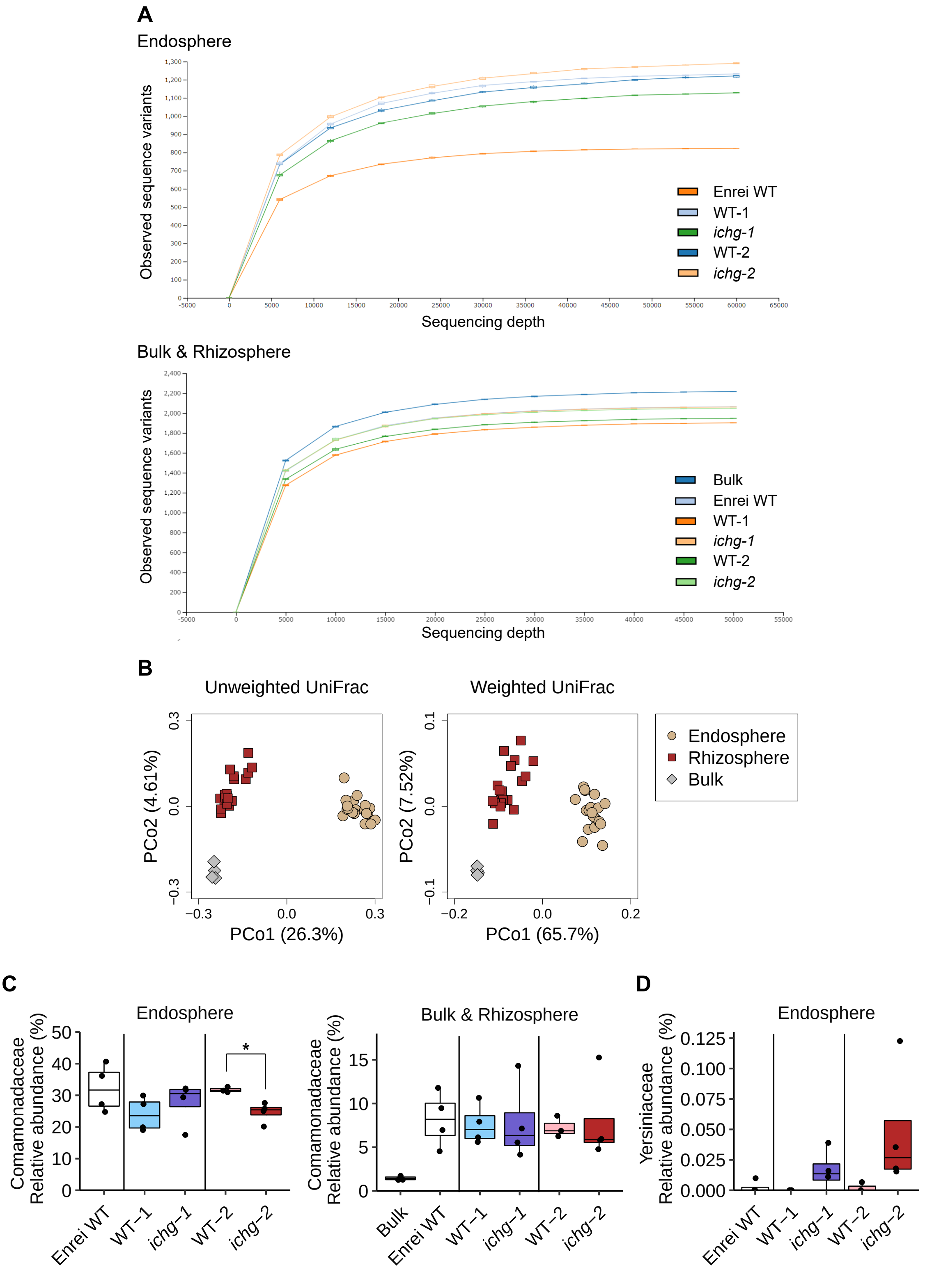


Supplementary Figure S1. Comparison of the bacterial communities from field-grown *ichg* mutants and WTs. (A) Rarefaction curves for the number of observed sequence variants in the endosphere and rhizosphere and bulk soil samples. (B) Unweighted and weighted UniFrac-based PCoA of the endosphere, rhizosphere, and bulk soil bacterial communities (endosphere and rhizosphere, n = 19; bulk soil, n = 4). Diamonds, squares, and circles indicate the bulk soil, rhizosphere, and endosphere, respectively. (C, D) Relative abundance of (C) *Comamonadaceae* and (D) *Yersiniaceae* in the endosphere and rhizosphere bacterial communities (WT-2, n = 3; the other genotypes, n = 4). The individual black dots indicate raw data points. The outliers were identified using the 1.5 × interquartile range rule. Student's two-tailed t-test was used for statistical analysis between the *ichg* mutant and WT allele from missense and nonsense mutants, respectively ($p < 0.05$). ICHG, isoflavone conjugate-hydrolyzing β-glucosidase; PCoA, principal coordinate analysis; WT, wild-type.

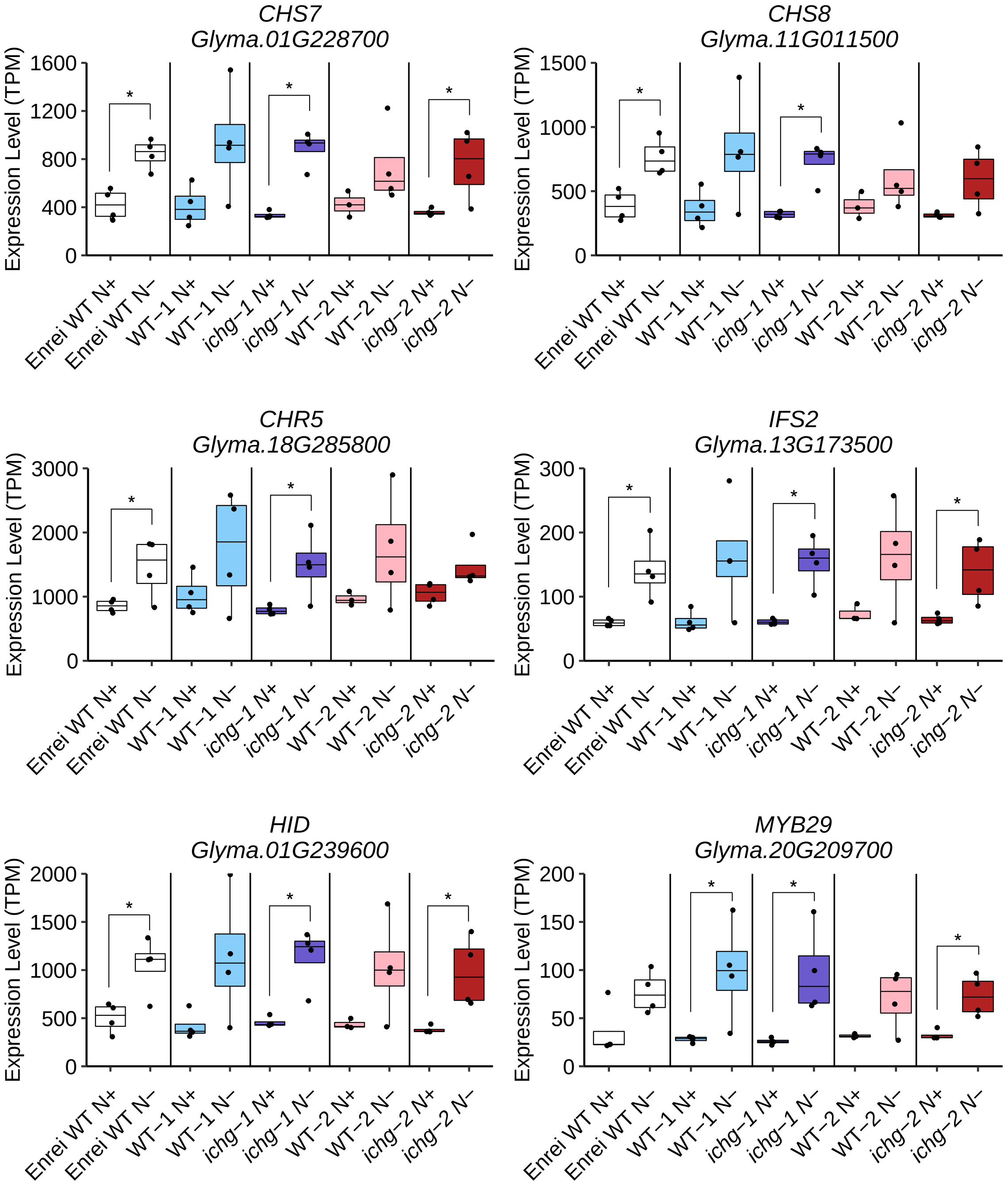


Supplementary Figure S2. mRNA levels of known genes up-regulated under nitrogen-deficient (N-) condition in roots of *ichg* mutants and WTs (WT-2 N+, n = 3; others, n = 4). The individual black dots indicate raw data points. The outliers were identified using the 1.5 × interquartile range rule. Student's two-tailed t-test was used for statistical analysis between nitrogen-sufficient (N+) and N- conditions in the same genotypes ($p < 0.05$). CHS, chalcone synthase; HID, 2-hydroxyisoflavanone dehydratase; ICHG, isoflavone conjugate-hydrolyzing β-glucosidase; IFS, isoflavone synthase; TPM, transcripts per million; WT, wild-type.

.

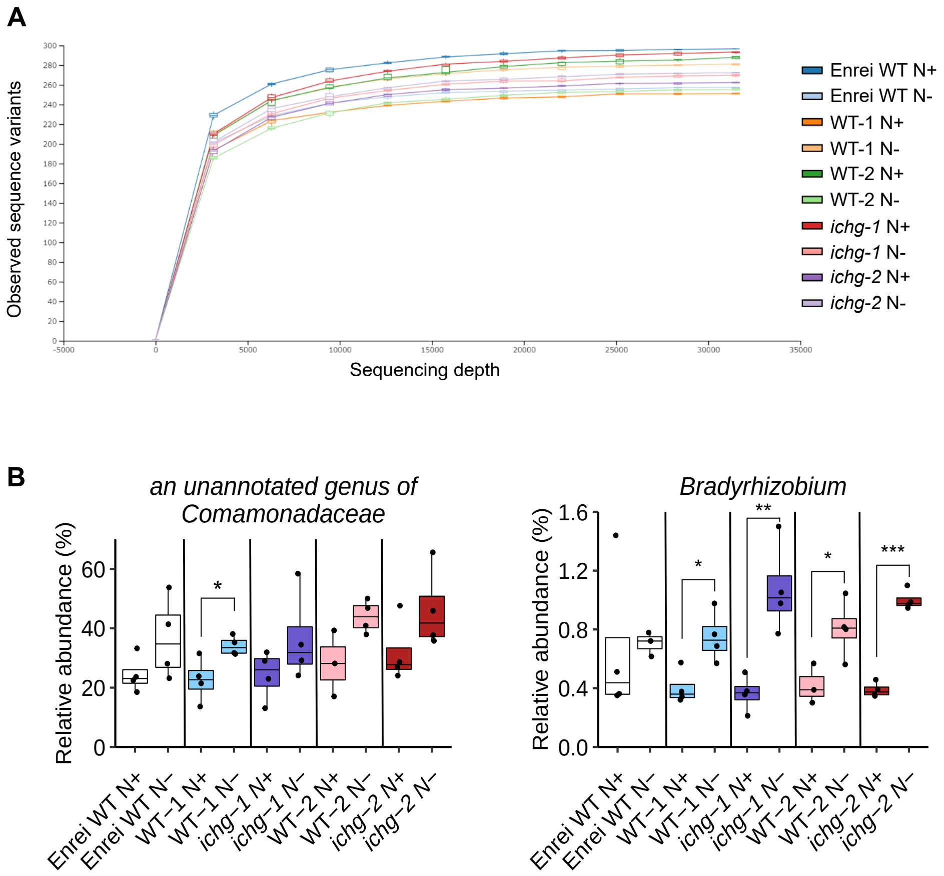


Supplementary Figure S3. Comparison of the root bacterial communities from *ichg* mutants and WTs cultivated under nitrogen-sufficient (N+) and -deficient (N-) conditions (WT-2 N+, n = 3; others, n = 4). (A) Rarefaction curves for the number of observed sequence variants. (B) Relative abundance of an unannotated genus of *Comamonadaceae* and *Bradyrhizobium*. The individual black dots indicate raw data points. The outliers were identified using the 1.5 × interquartile range rule. For *Bradyrhizobium*, one replicate in Enrei WT N- is excluded in the boxplot due to its extremely high abundance compared to other samples. Student's two-tailed t-test was used for statistical analysis between N+ and N- conditions in the same genotypes (* $p < 0.05$, ** $p < 0.01$, *** $p < 0.001$). ICHG, isoflavone conjugate-hydrolyzing β-glucosidase; WT, wild-type.

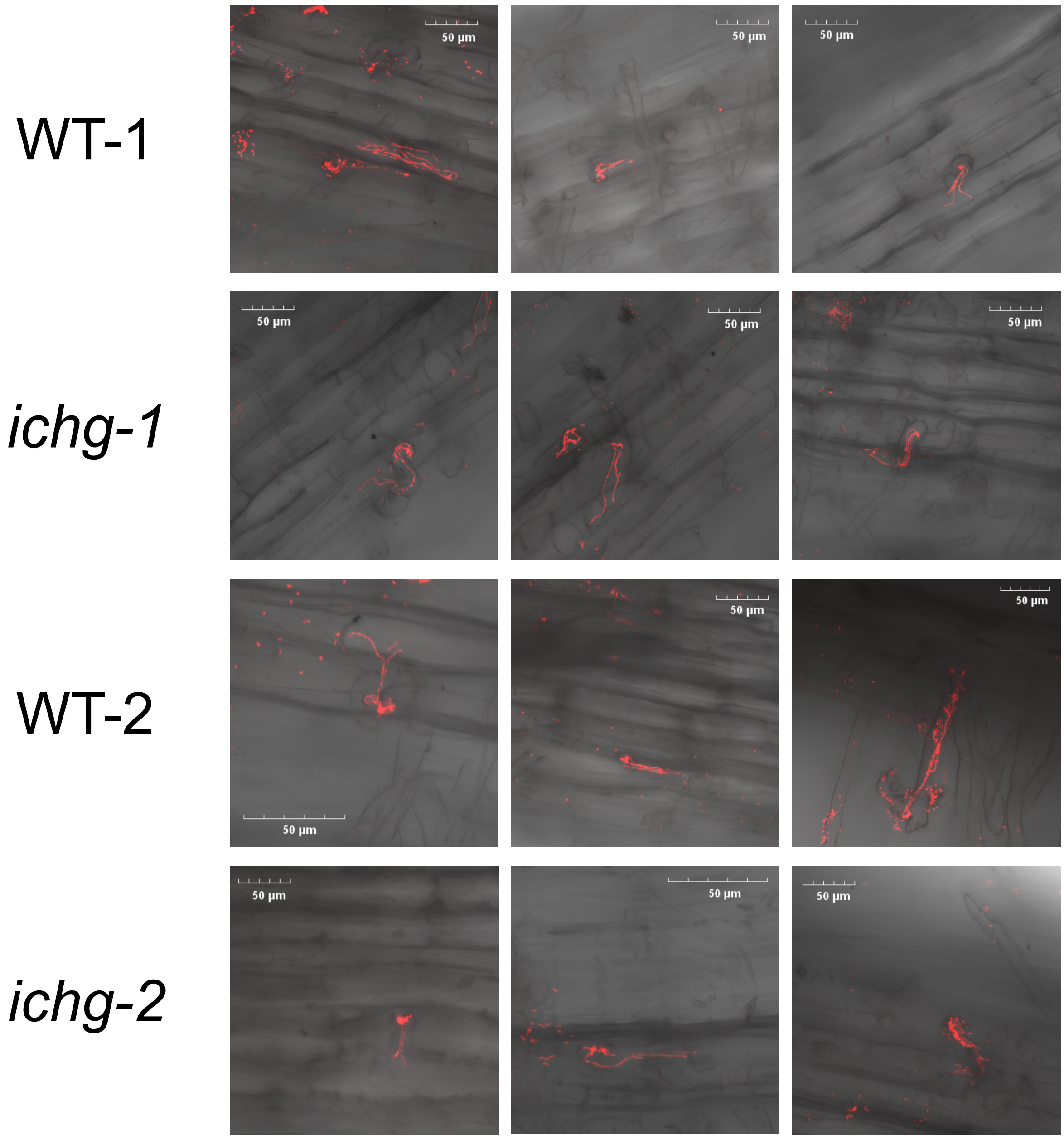


Supplementary Figure S4. Fluorescent images of the DsRed-labeled rhizobial infection threads on soybean roots. *Bradyrhizobium diazoefficiens* USDA 110 was inoculated onto 5-day-old soybean seedlings under sterile conditions and infection threads on the roots were observed after 7 days. Three representative shots of different infection threads for each genotype are shown. ICHG, isoflavone conjugate-hydrolyzing β-glucosidase; WT, wild-type.

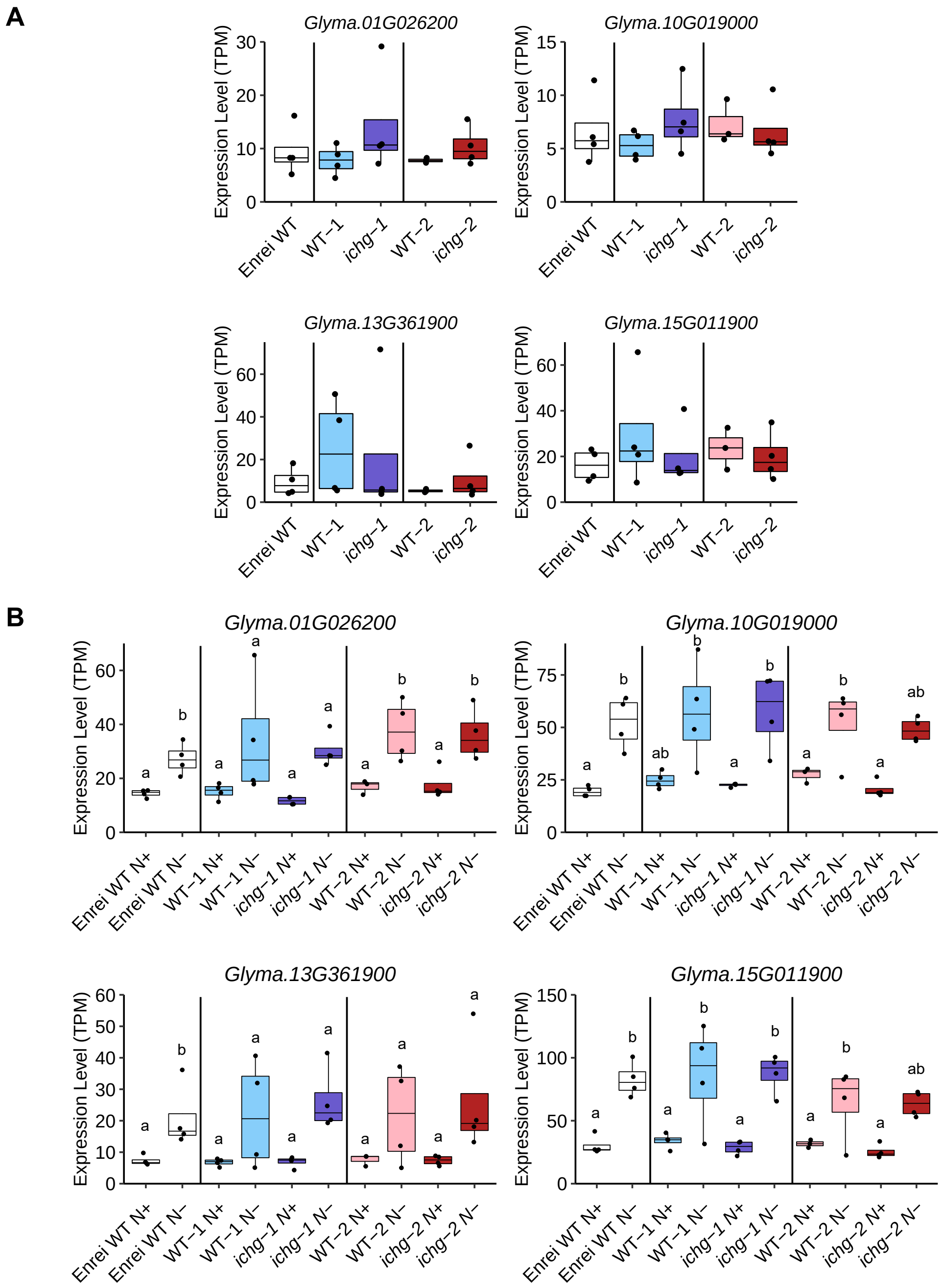


Supplementary Figure S5. mRNA levels of transporter genes in roots of *ichg* mutants and WTs cultivated in (A) field and (B) greenhouse. These transporter genes were co-expressed diurnally (r > 0.8) with isoflavone biosynthetic genes (Matsuda et al., 2020). *Glyma.01G026200*, a multidrug and toxin extrusion transporter; *Glyma.10G019000*, a ATP-binding cassette (ABC) transporter C subfamily; *Glyma.13G361900* and *Glyma.15G011900*, ABC transporter G subfamily. The individual black dots indicate raw data points. The outliers were identified using the 1.5 × interquartile range rule. (A) Student's two-tailed t-test was used for statistical analysis between nitrogen-sufficient (N+) and nitrogen-deficient (N-) conditions in the same genotypes (WT-2, n = 3; others, n = 4) ($p < 0.05$). (B) Tukey's HSD test was used for statistical analysis among Enrei WT, the *ichg* mutant and WT allele from missense and nonsense mutants, respectively (WT-2 N+, n = 3; others, n = 4) ($p < 0.05$). ICHG, isoflavone conjugate-hydrolyzing β-glucosidase; TPM, transcripts per million; WT, wild-type.

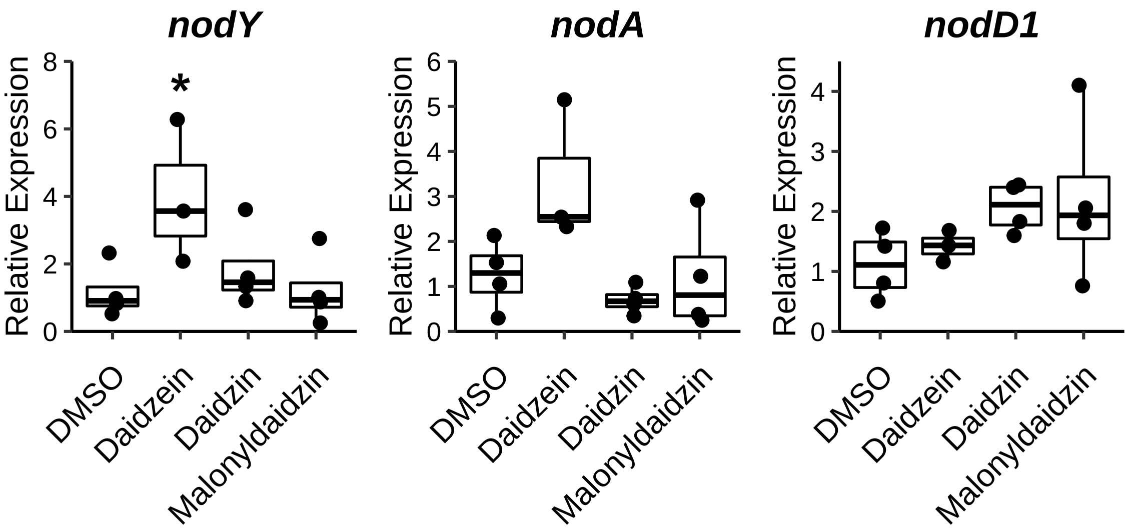


Supplementary Figure S6. Relative expression levels of *nod* genes of *Bradyrhizobium diazoefficiens* USDA110 inducted by isoflavone standard chemicals. Relative gene expression was calculated with the Pfaffl method using *ParA* as a reference gene. Dunnett's test was used for statistical analysis on daidzein, daidzin, and malonyldaidzin-treated samples compared to DMSO-treated sample ($p < 0.05$). DMSO, dimethyl sulfoxide.
