## Supplemental Tables for "Apoplast-localized β-Glucosidase Elevates Isoflavone Accumulation in the Soybean Rhizosphere"

Supplementary Table S1. PERMANOVA results for the first two PCs in the principal component analysis of transcriptome of field-grown *ichg* mutants and wild-types (WTs).

Leaf

| Group 1 | Group 2 | Sample size | Permutations | p-value | q-value |
|---|---|---|---|---|---|
| Enrei WT | *ichg-2* | 8 | 1000 | 0.311688312 | 0.346320346 |
| Enrei WT | *ichg-1* | 8 | 1000 | 0.217782218 | 0.272227772 |
| Enrei WT | WT-1 | 8 | 1000 | 0.026973027 | 0.093240093 |
| Enrei WT | WT-2 | 7 | 1000 | 0.174825175 | 0.24975025 |
| WT-1 | *ichg-1* | 8 | 1000 | 0.065934066 | 0.164835165 |
| WT-1 | WT-2 | 7 | 1000 | 0.027972028 | 0.093240093 |
| WT-1 | *ichg-2* | 8 | 1000 | 0.018981019 | 0.093240093 |
| WT-2 | *ichg-2* | 7 | 1000 | 0.822177822 | 0.822177822 |
| WT-2 | *ichg-1* | 7 | 1000 | 0.14985015 | 0.24975025 |
| *ichg-2* | *ichg-1* | 8 | 1000 | 0.10989011 | 0.21978022 |

Root

| Group 1 | Group 2 | Sample size | Permutations | p-value | q-value |
|---|---|---|---|---|---|
| Enrei WT | *ichg-2* | 8 | 1000 | 0.843156843 | 0.852147852 |
| Enrei WT | *ichg-1* | 8 | 1000 | 0.756243756 | 0.828060828 |
| Enrei WT | WT-1 | 8 | 1000 | 0.507492507 | 0.828060828 |
| Enrei WT | WT-2 | 7 | 1000 | 0.655344655 | 0.828060828 |
| WT-1 | *ichg-1* | 8 | 1000 | 0.053946054 | 0.569430569 |
| WT-1 | WT-2 | 7 | 1000 | 0.410589411 | 0.828060828 |
| WT-1 | *ichg-2* | 8 | 1000 | 0.26973027 | 0.828060828 |
| WT-2 | *ichg-2* | 7 | 1000 | 0.739260739 | 0.828060828 |
| WT-2 | *ichg-1* | 7 | 1000 | 0.548451548 | 0.828060828 |
| *ichg-2* | *ichg-1* | 8 | 1000 | 0.651348651 | 0.828060828 |

Supplementary Table S2. Number of DEGs in the *ichg* mutants when compared with the *ICHG* WTs (false discovery rate < 0.01).

| Organ | *ICHG* WT | *ichg* mutant | Up-regulated DEGs in the *ichg* mutant | Down-regulated DEGs in the *ichg* mutant |
|---|---|---|---|---|
| Leaf | WT-1 | *ichg-1* | 34 | 152 |
| Leaf | WT-2 | *ichg-2* | 3 | 2 |
| Root | WT-1 | *ichg-1* | 128 | 177 |
| Root | WT-2 | *ichg-2* | 13 | 7 |

Supplementary Table S3. Unweighted and weighted UniFrac-based PERMANOVA results for the endosphere, rhizosphere, and bulk soil samples.

Unweighted

| Group 1 | Group 2 | Sample size | Permutations | pseudo-F | p-value | q-value |
|---|---|---|---|---|---|---|
| Bulk | Endosphere | 23 | 999 | 7.052272014 | 0.001 | 0.001 |
| Bulk | Rhizosphere | 23 | 999 | 2.32333371 | 0.001 | 0.001 |
| Endosphere | Rhizosphere | 38 | 999 | 12.6373636 | 0.001 | 0.001 |

Weighted

| Group 1 | Group 2 | Sample size | Permutations | pseudo-F | p-value | q-value |
|---|---|---|---|---|---|---|
| Bulk | Endosphere | 23 | 999 | 40.64219989 | 0.001 | 0.001 |
| Bulk | Rhizosphere | 23 | 999 | 11.9195821 | 0.001 | 0.001 |
| Endosphere | Rhizosphere | 38 | 999 | 61.44367948 | 0.001 | 0.001 |

Supplementary Table S4. Unweighted and weighted UniFrac-based PERMANOVA results for the endosphere bacterial communities of each sample.

Unweighted

| Group 1 | Group 2 | Sample size | Permutations | pseudo-F | p-value | q-value |
|---|---|---|---|---|---|---|
| WT-2 | Enrei WT | 7 | 999 | 1.19067282 | 0.067 | 0.243333333 |
| WT-2 | *ichg-2* | 7 | 999 | 0.990621839 | 0.517 | 0.626666667 |
| WT-2 | *ichg-1* | 7 | 999 | 1.014015271 | 0.368 | 0.626666667 |
| WT-1 | WT-2 | 7 | 999 | 1.022083139 | 0.439 | 0.626666667 |
| WT-1 | Enrei WT | 8 | 999 | 1.133210805 | 0.139 | 0.3475 |
| WT-1 | *ichg-2* | 8 | 999 | 0.981791595 | 0.515 | 0.626666667 |
| WT-1 | *ichg-1* | 8 | 999 | 0.954280478 | 0.655 | 0.655 |
| Enrei WT | *ichg-2* | 8 | 999 | 1.20599393 | 0.073 | 0.243333333 |
| Enrei WT | *ichg-1* | 8 | 999 | 1.161373059 | 0.02 | 0.2 |
| *ichg-1* | *ichg-2* | 8 | 999 | 0.980286619 | 0.564 | 0.626666667 |

Weighted

| Group 1 | Group 2 | Sample size | Permutations | pseudo-F | p-value | q-value |
|---|---|---|---|---|---|---|
| WT-2 | Enrei WT | 7 | 999 | 1.681407672 | 0.132 | 0.626666667 |
| WT-2 | *ichg-2* | 7 | 999 | 1.226610089 | 0.323 | 0.626666667 |
| WT-2 | *ichg-1* | 7 | 999 | 0.767311707 | 0.761 | 0.784 |
| WT-1 | WT-2 | 7 | 999 | 1.433973604 | 0.21 | 0.626666667 |
| WT-1 | Enrei WT | 8 | 999 | 1.731430592 | 0.087 | 0.626666667 |
| WT-1 | *ichg-2* | 8 | 999 | 1.11528388 | 0.459 | 0.655714286 |
| WT-1 | *ichg-1* | 8 | 999 | 0.948288111 | 0.584 | 0.73 |
| Enrei WT | *ichg-2* | 8 | 999 | 1.53044306 | 0.333 | 0.626666667 |
| Enrei WT | *ichg-1* | 8 | 999 | 1.117362287 | 0.376 | 0.626666667 |
| *ichg-1* | *ichg-2* | 8 | 999 | 0.668559233 | 0.784 | 0.784 |

Supplementary Table S5. Unweighted and weighted UniFrac-based PERMANOVA results for the rhizosphere bacterial communities of each sample.

Unweighted

| Group 1 | Group 2 | Sample size | Permutations | pseudo-F | p-value | q-value |
|---|---|---|---|---|---|---|
| Enrei WT | WT-1 | 8 | 999 | 0.997781198 | 0.486 | 0.6625 |
| Enrei WT | WT-2 | 7 | 999 | 0.976544451 | 0.65 | 0.722222222 |
| Enrei WT | *ichg-1* | 8 | 999 | 1.070422973 | 0.216 | 0.618 |
| Enrei WT | *ichg-2* | 8 | 999 | 0.989645466 | 0.53 | 0.6625 |
| WT-1 | WT-2 | 7 | 999 | 1.103106036 | 0.096 | 0.48 |
| WT-1 | *ichg-1* | 8 | 999 | 0.997305777 | 0.409 | 0.6625 |
| WT-1 | *ichg-2* | 8 | 999 | 1.100591641 | 0.048 | 0.48 |
| WT-2 | *ichg-1* | 7 | 999 | 1.028859065 | 0.309 | 0.618 |
| WT-2 | *ichg-2* | 7 | 999 | 0.925190091 | 1 | 1 |
| ichg-1 | *ichg-2* | 8 | 999 | 1.04884679 | 0.256 | 0.618 |

Weighted

| Group 1 | Group 2 | Sample size | Permutations | pseudo-F | p-value | q-value |
|---|---|---|---|---|---|---|
| Enrei WT | WT-1 | 8 | 999 | 2.069726247 | 0.09 | 0.313333333 |
| Enrei WT | WT-2 | 7 | 999 | 1.505925591 | 0.171 | 0.4275 |
| Enrei WT | *ichg-1* | 8 | 999 | 1.512148334 | 0.219 | 0.438 |
| Enrei WT | *ichg-2* | 8 | 999 | 0.906085479 | 0.477 | 0.605 |
| WT-1 | WT-2 | 7 | 999 | 2.585206229 | 0.061 | 0.313333333 |
| WT-1 | *ichg-1* | 8 | 999 | 0.785509688 | 0.678 | 0.678 |
| WT-1 | *ichg-2* | 8 | 999 | 1.834221765 | 0.094 | 0.313333333 |
| WT-2 | *ichg-1* | 7 | 999 | 0.964009653 | 0.484 | 0.605 |
| WT-2 | *ichg-2* | 7 | 999 | 0.87897776 | 0.61 | 0.677777778 |
| ichg-1 | *ichg-2* | 8 | 999 | 0.832340882 | 0.476 | 0.605 |

Supplementary Table S6. Number of enriched and depleted bacterial families in the *ichg* mutants when compared with *ICHG* WTs using LEfSe method (LDA score >2.0).

| Region | *ICHG* WT | *ichg* mutant | Enriched families in the *ichg* mutant | Depleted families in the *ichg* mutant |
|---|---|---|---|---|
| Endosphere | WT-1 | *ichg-1* | 2 | 1 |
| Endosphere | WT-2 | *ichg-2* | 6 | 5 |
| Rhizosphere | WT-1 | *ichg-1* | 2 | 4 |
| Rhizosphere | WT-2 | *ichg-2* | 2 | 3 |

Supplementary Table S7. PERMANOVA results for the first two PCs in the principal component analysis of the root transcriptome of *ichg* mutants and WTs cultivated under nitrogen-sufficient (N+) and nitrogen-deficient (N-) conditions.

Nutrition

| Group 1 | Group 2 | Sample size | Permutations | p-value | q-value |
|---|---|---|---|---|---|
| N+ | N- | 39 | 1000 | 0.000999001 | 0.004995005 |

Genotype

| Group 1 | Group 2 | Sample size | Permutations | p-value | q-value |
|---|---|---|---|---|---|
| Enrei WT | *ichg-2* | 16 | 1000 | 0.000999001 | 0.004995005 |
| Enrei WT | *ichg-1* | 16 | 1000 | 0.904095904 | 0.968674183 |
| Enrei WT | WT-1 | 16 | 1000 | 0.398601399 | 0.543547362 |
| Enrei WT | WT-2 | 15 | 1000 | 0.033966034 | 0.101898102 |
| WT-1 | *ichg-1* | 16 | 1000 | 0.599400599 | 0.697379544 |
| WT-1 | WT-2 | 15 | 1000 | 0.380619381 | 0.543547362 |
| WT-1 | *ichg-2* | 16 | 1000 | 0.010989011 | 0.041208791 |
| WT-2 | *ichg-2* | 15 | 1000 | 0.20979021 | 0.34965035 |
| WT-2 | *ichg-1* | 15 | 1000 | 0.068931069 | 0.147709433 |
| *ichg-2* | *ichg-1* | 16 | 1000 | 0.000999001 | 0.004995005 |

Supplementary Table S8. Unweighted and weighted UniFrac-based PERMANOVA results for endosphere samples cultivated under N+ and N- conditions.

Unweighted

| Group 1 | Group 2 | Sample size | Permutations | pseudo-F | p-value | q-value |
|---|---|---|---|---|---|---|
| N+ | N- | 39 | 999 | 2.790366807 | 0.001 | 0.001 |

Weighted

| Group 1 | Group 2 | Sample size | Permutations | pseudo-F | p-value | q-value |
|---|---|---|---|---|---|---|
| N+ | N- | 39 | 999 | 7.326676211 | 0.001 | 0.001 |

Supplementary Table S9. Unweighted and weighted UniFrac-based PERMANOVA results for the endosphere bacterial communities under N+ condition.

Unweighted

| Group 1 | Group 2 | Sample size | Permutations | pseudo-F | p-value | q-value |
|---|---|---|---|---|---|---|
| Enrei WT | WT-1 | 8 | 999 | 1.414672496 | 0.054 | 0.27 |
| Enrei WT | WT-2 | 7 | 999 | 0.903943995 | 0.622 | 0.753333333 |
| Enrei WT | ichg-1 | 8 | 999 | 0.932885329 | 0.402 | 0.67 |
| Enrei WT | ichg-2 | 8 | 999 | 1.017145009 | 0.361 | 0.67 |
| WT-1 | WT-2 | 7 | 999 | 1.550283334 | 0.023 | 0.23 |
| WT-1 | ichg-1 | 8 | 999 | 0.919879707 | 0.678 | 0.753333333 |
| WT-1 | ichg-2 | 8 | 999 | 1.109473172 | 0.248 | 0.62 |
| WT-2 | ichg-1 | 7 | 999 | 1.180220121 | 0.211 | 0.62 |
| WT-2 | ichg-2 | 7 | 999 | 0.887504133 | 0.826 | 0.826 |
| ichg-1 | ichg-2 | 8 | 999 | 0.957923923 | 0.594 | 0.753333333 |

Weighted

| Group 1 | Group 2 | Sample size | Permutations | pseudo-F | p-value | q-value |
|---|---|---|---|---|---|---|
| Enrei WT | WT-1 | 8 | 999 | 1.974898275 | 0.107 | 0.32 |
| Enrei WT | WT-2 | 7 | 999 | 1.429977076 | 0.259 | 0.431666667 |
| Enrei WT | ichg-1 | 8 | 999 | 1.671815295 | 0.128 | 0.32 |
| Enrei WT | ichg-2 | 8 | 999 | 2.28947231 | 0.059 | 0.32 |
| WT-1 | WT-2 | 7 | 999 | 0.844902314 | 0.599 | 0.678888889 |
| WT-1 | ichg-1 | 8 | 999 | 1.113880158 | 0.447 | 0.638571429 |
| WT-1 | ichg-2 | 8 | 999 | 0.738036159 | 0.611 | 0.678888889 |
| WT-2 | ichg-1 | 7 | 999 | 1.888484413 | 0.189 | 0.378 |
| WT-2 | ichg-2 | 7 | 999 | 0.417414299 | 0.91 | 0.91 |
| ichg-1 | ichg-2 | 8 | 999 | 1.908717359 | 0.111 | 0.32 |

Supplementary Table S10. Unweighted and weighted UniFrac-based PERMANOVA results for the endosphere bacterial communities under N- condition.

Unweighted

| Group 1 | Group 2 | Sample size | Permutations | pseudo-F | p-value | q-value |
|---|---|---|---|---|---|---|
| Enrei WT | WT-1 | 8 | 999 | 1.013508093 | 0.437 | 0.785 |
| Enrei WT | WT-2 | 8 | 999 | 1.033285682 | 0.323 | 0.785 |
| Enrei WT | ichg-1 | 8 | 999 | 1.038772072 | 0.449 | 0.785 |
| Enrei WT | ichg-2 | 8 | 999 | 0.85366061 | 0.724 | 0.785 |
| WT-1 | WT-2 | 8 | 999 | 0.937513226 | 0.591 | 0.785 |
| WT-1 | ichg-1 | 8 | 999 | 1.130811306 | 0.267 | 0.785 |
| WT-1 | ichg-2 | 8 | 999 | 0.824261874 | 0.738 | 0.785 |
| WT-2 | ichg-1 | 8 | 999 | 0.874212733 | 0.785 | 0.785 |
| WT-2 | ichg-2 | 8 | 999 | 0.9306205 | 0.69 | 0.785 |
| ichg-1 | ichg-2 | 8 | 999 | 1.083004285 | 0.313 | 0.785 |

Weighted

| Group 1 | Group 2 | Sample size | Permutations | pseudo-F | p-value | q-value |
|---|---|---|---|---|---|---|
| Enrei WT | WT-1 | 8 | 999 | 1.024364169 | 0.528 | 0.686 |
| Enrei WT | WT-2 | 8 | 999 | 3.405327661 | 0.078 | 0.39 |
| Enrei WT | ichg-1 | 8 | 999 | 0.532988113 | 0.686 | 0.686 |
| Enrei WT | ichg-2 | 8 | 999 | 1.404781732 | 0.318 | 0.608333333 |
| WT-1 | WT-2 | 8 | 999 | 5.072350694 | 0.021 | 0.21 |
| WT-1 | ichg-1 | 8 | 999 | 0.900576882 | 0.557 | 0.686 |
| WT-1 | ichg-2 | 8 | 999 | 1.385909399 | 0.336 | 0.608333333 |
| WT-2 | ichg-1 | 8 | 999 | 2.799629279 | 0.149 | 0.496666667 |
| WT-2 | ichg-2 | 8 | 999 | 0.615230702 | 0.65 | 0.686 |
| ichg-1 | ichg-2 | 8 | 999 | 1.163412514 | 0.365 | 0.608333333 |
